# Transcriptional changes are regulated by metabolic pathway dynamics but decoupled from protein levels

**DOI:** 10.1101/833921

**Authors:** Jack E. Feltham, Shidong Xi, Struan C. Murray, Meredith Wouters, Julian Urdiain-Arraiza, Raphael Heilig, Charlotte George, Anna F. Townley, Emile Roberts, Benedikt M. Kessler, Sabrina Liberatori, Philip D. Charles, Andrew Angel, Roman Fischer, Jane Mellor

## Abstract

Transcription is necessary for the synthesis of new proteins, often leading to the assumption that changes in transcript levels lead to changes in protein levels which directly impact a cell’s phenotype. Using a synchronized biological rhythm, we show that despite genome-wide partitioning of transcription, transcripts and translation levels into two phase-shifted expression clusters related to metabolism, detectable protein levels remain constant over time. This disconnect between cycling translation and constant protein levels can be explained by slow protein turnover rates, with overall protein levels maintained by low level pulses of new protein synthesis. Instead, rhythmic post-translational regulation of the activities of different proteins, influenced by the metabolic state of the cells, appears to be key to coordinating the physiology of the biological rhythm with cycling transcription. Thus, transcriptional and translational cycling reflects, rather than drives, metabolic and biosynthetic changes during biological rhythms. We propose that transcriptional changes are often the consequence, rather than the cause, of changes in cellular physiology and that caution is needed when inferring the activity of biological processes from transcript data.

- Changes in protein levels do not explain the changing states of a biological rhythm
- Slow protein turnover rates decouple proteins levels from a rhythmic transcriptome
- Metabolites determine protein activity via rhythmic post-translational modifications
- Cycling protein activity explains rhythmic transcription and ribosome biogenesis
- A cycling transcriptome is a consequence, not a cause, of physiological changes

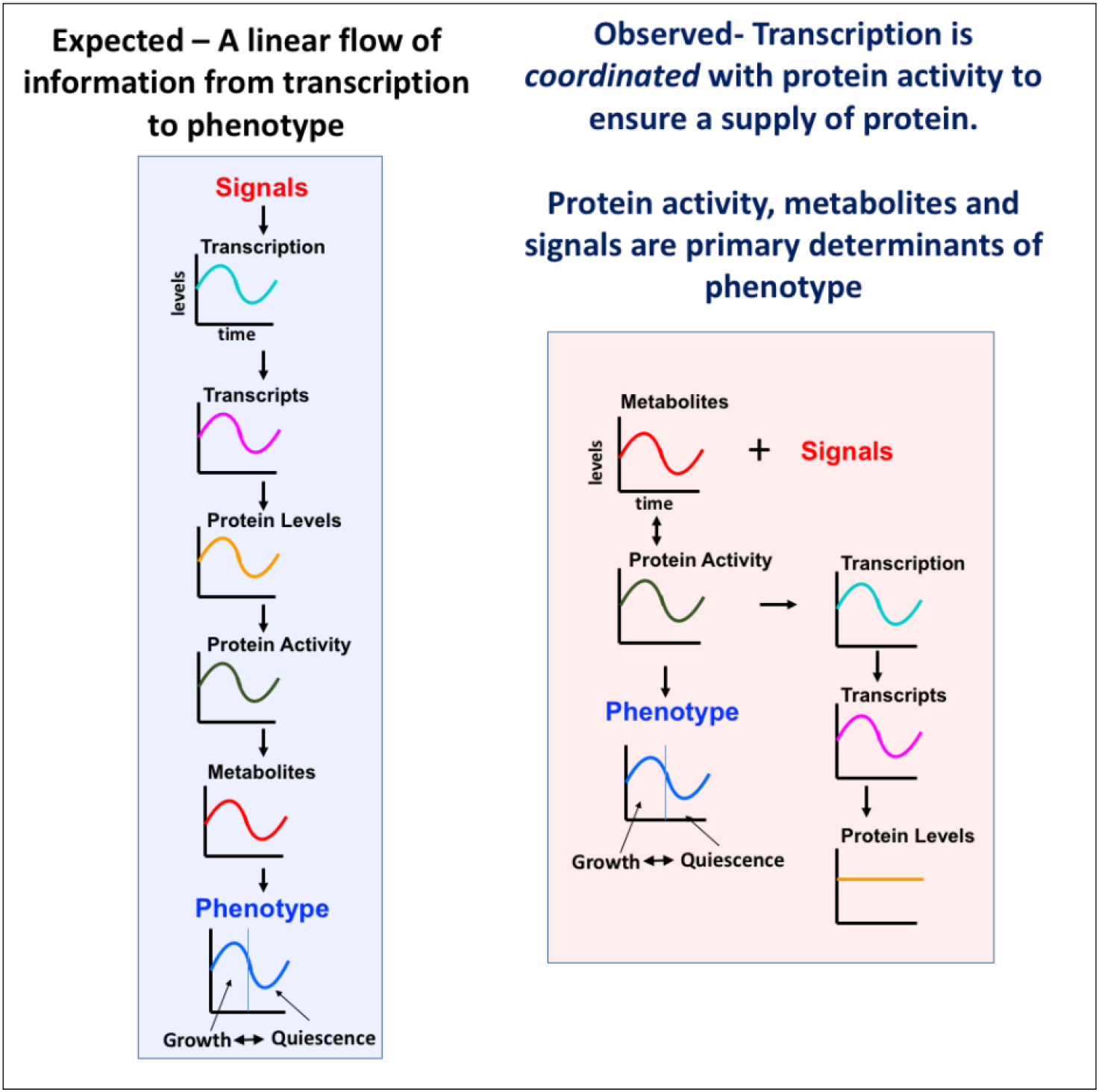

## Introduction

It is universally accepted that RNA synthesis is required for protein production in eukaryotic cells. In response to developmental cues and stress-responses, transcriptional induction of protein-coding genes or stabilization of their transcripts drives accumulation of their protein products. The proteins in turn remodel cellular physiology in response to the stimulus (Banerji et al., 1984; Storti et al., 1980; Tissieres et al., 1974). The ubiquity of this underlying axis of information-transfer can easily draw one to the conclusion that whenever transcript abundances change, protein abundance and cellular physiology will follow suit. However, recent studies are beginning to question this assumption (Caster et al., 2016; Liu et al., 2016; Noya et al., 2019; Ray and Reddy, 2016). In the proteomics field, the lack of correlation between the transcriptome and the proteome has been known for some years but remains largely unexplained (Vogel and Marcotte, 2012). This has important implications for research because RNA-based assays are so widely used to infer the behaviour of proteins and associated phenotypes.

Cycling transcripts are a universal feature of eukaryotic biological rhythms (Benegiamo et al., 2016; Mellor, 2016) and when these rhythms are synchronized, they provide an ideal system in which to assess how changes in the levels of both nascent and stable transcripts affect the proteome. Based on oscillating levels of transcripts, many have argued that cycling levels of gene products might act to regulate biological rhythms. These include circadian rhythms, which are fundamentally metabolic oscillations with additional time keeping mechanisms, and the respiratory and metabolic oscillations in yeast, also known as the yeast metabolic cycle (YMC) (Klevecz et al., 2004; Kuang et al., 2014; Takahashi, 2017; Tu et al., 2005). In the YMC, yeast show synchronized phases of growth and quiescence, linked to metabolism and phases of high and low oxygen consumption (HOC and LOC) (Causton et al., 2015; Klevecz et al., 2004; Kuang et al., 2014; Mellor, 2016; Tu et al., 2005; Tu and McKnight, 2006). Here we report a multi-omic analysis (Sánchez-Gaya et al., 2018) of the YMC, from the nascent transcriptome through to the proteome. Despite cycling transcripts and translation, detectable protein levels remain constant over the rhythm. The disconnect between cycling transcripts and constant protein levels can be explained by slow protein turnover rates, with overall protein levels maintained by low level pulses of new protein synthesis. Thus the physiology of the rhythm cannot be driven by cycling proteins. Instead, post-translational regulation of protein activity, influenced by the metabolic state of the cells, appears to be key to maintaining the biological rhythm and cycling transcription. We propose, contrary to the prevailing view, that transcriptional changes are often the consequence, rather than the cause, of changes in cellular physiology.

## Results

### The cycling transcriptome forms two phase-shifted clusters related to metabolism

To assess levels of nascent and steady-state transcripts, 11 calibrated NET-Seq (Churchman and Weissman, 2011; Fischl et al., 2017) libraries were generated alongside 22 calibrated polyA-primed RNASeq libraries, throughout the synchronised biological rhythm of *S.cerevisiae.* The temporal position of the cells in the rhythm, and their synchronous entry into the cell cycle, was monitored using the change in dissolved O2 in the growth medium (Figure 1A,B, EV1A, EV2A,B), partitioning cycling into phases of high and low oxygen consumption (HOC, LOC). These phases have been associated with metabolites and cellular processes mediating growth and quiescence, respectively. Moreover, the LOC phase appears to be the default state, as the frequency with which cells enter the fixed-length HOC is determined by nutrient availability (Mellor, 2016). The NET-Seq libraries (Figure EV1B-D) were log_2_ fold-change vs median (LFCM) transformed and divided into three clusters of nascent transcripts (T), as recommended by the gap-statistic (Figure EV1E), using the k-means++ algorithm, and plotted as metagenes (Figure 1C). This revealed one non-cycling cluster (NC-T) and two cycling clusters with large log_2_ foldchanges that coincided with the HOC phase of the rhythm. One cluster exhibited an oscillation with maximum levels of nascent transcripts during the LOC phase, anti-phase to the dissolved O2 levels (cluster LOC-T, 805 genes, mean NET-Seq range 2.7-fold). The second cluster containing 848 genes, peaked in expression during the HOC phase (cluster HOC-T, mean NET-Seq range 6.5-fold), before dropping sharply at the HOC/LOC boundary then returning to basal expression levels. Considering the possibility that lowly transcribed genes would be differently amenable to changes in levels of nascent transcript production to highly transcribed ones, we plotted the absolute log_2_-NET-Seq signals by cluster as octiles (Figure EV1F). Surprisingly, all clusters showed similar variation in nascent transcript levels and, although HOC-T and LOC-T both cycled over several orders of magnitude, substantial levels of transcription were observed even at the lowest points. In addition, the three clusters each have similar median expression levels (Figure EV1G).

**Figure 1.**
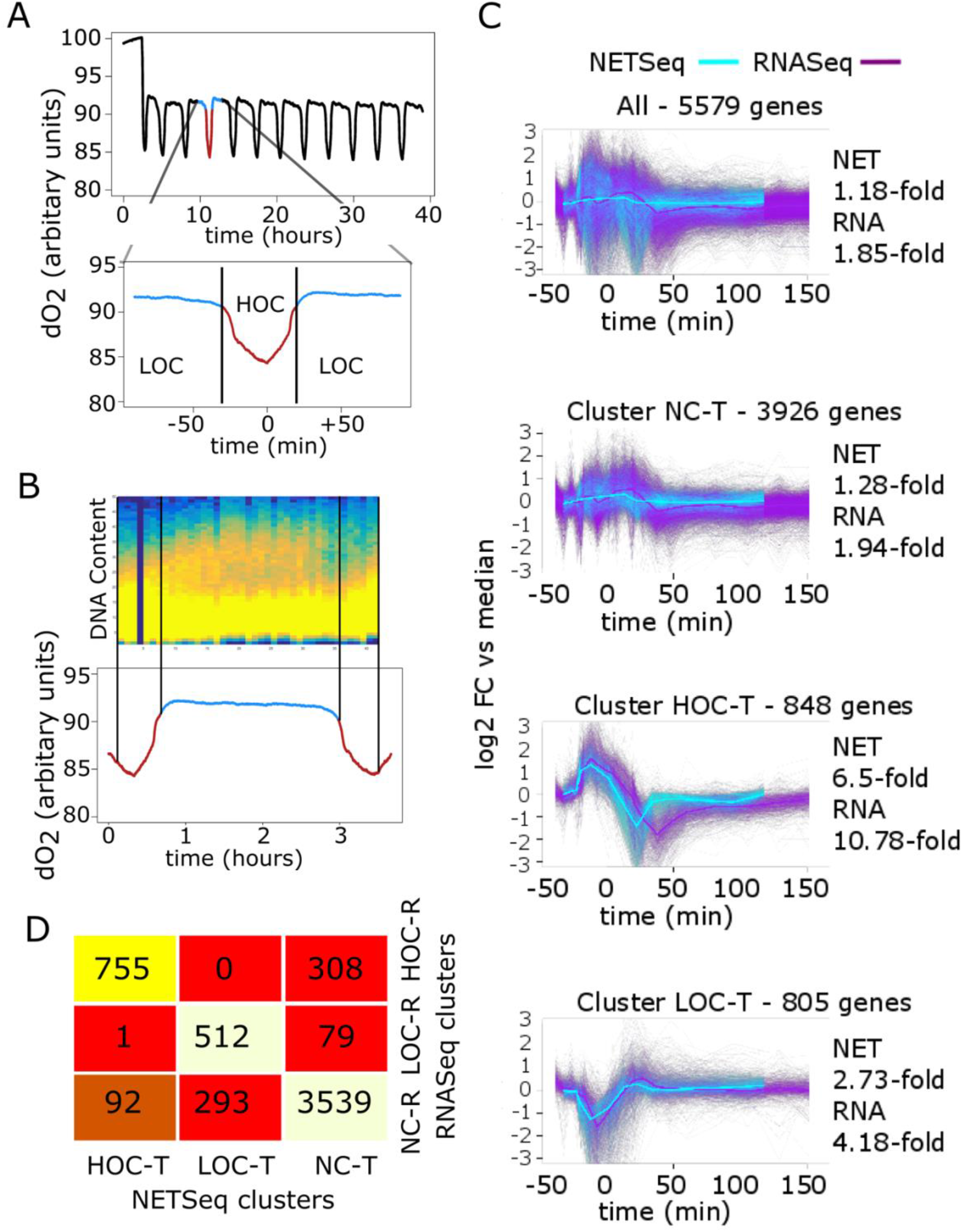
The cycling transcriptome of the YMC is primarily driven by transcriptional oscillations. **(A)** Alternative phases of high and low oxygen consumption in the YMC assessed by levels of dissolved oxygen (dO_2_) in a synchronised culture. These profiles allow sampling to be coordinated between experiments as the cycle length is highly reproducible. (**B**) FACS analysis of strains stained with propidium iodine revealing the DNA content and the period of S phase over time. About 10% of cells enter S phase during the HOC phase. (**C**) 11 NETSeq and 22 polyA-primed RNASeq samples were taken at the time indicated (see EV Table 1 for details; n=1 for each complete data set shown here; n=3 for RNASeq data sets in total (22 time points; 11 time points to match with the NETSeq data and the 20 samples for the polysome profiling input) from *CENPK113-7D rpb3-FLAG::KANMX6* cells cycling at a dilution rate of 0.082 hr^-1^. Counts tables were generated from both datasets for all *S. cerevisiae* genes and for each gene and dataset, a log_2_ fold-change vs median (LFCM) transformation was applied. Gap-Statistic analysis was applied to the transformed NETSeq data to determine the correct number of clusters into which to divide genes using the k-means++ algorithm. Transformed NETSeq and RNASeq data was divided into the same three NETSeq clusters and plotted over time alongside all genes. (**D**) For both datasets, the transformed data was divided into three clusters using the k-means++ algorithm and the cluster:cluster overlap was plotted as a heatmap visualizing the self-correlation matrix of the three clusters for RNA-samples derived by NETSeq (T) or RNASeq (R). Warm colours (yellow/white) represent high correlations and similarities. For details see Figures EV1 and EV2.

Next, we asked how steady-state RNA levels related to levels of nascent transcripts (Figure 1C, EV2). Again, genes were clustered by log_2_ fold-change vs median (LFCM) using the k-means++ algorithm. Gap-statistic analysis indicated that the data was best divided into either one or three clusters (Figure EV2C). The three cluster division was chosen for analysis in order to better reflect the analyses performed in previous studies which concluded that majority of the transcriptome oscillates in 3-7 clusters, arguing that cycling levels of gene products regulate the metabolic and biosynthetic pathways during the cycle (Klevecz et al., 2004; Kuang et al., 2014; Tu et al., 2005). Reassuringly, the three steadystate RNA (R) clusters (Figure EV2D,E) bore a close relationship with the NETSeq (T) clusters, falling into the same broad categories (NC, HOC and LOC), yet some differences can be observed, suggesting post-transcriptional control of steady-state transcript levels (Figure 1C,D, EV2F,G). For example, in Figure 1D, 308 members of the HOC-R RNASeq cluster are in the NC-T NETSeq cluster, suggesting that they are regulated at the post-transcriptional level, through differential RNA stability, as described for mitochondrial RPG transcripts (Kuang et al., 2014; Lee and Tu, 2015; Tu et al., 2005). Similarly, the NC-R and LOC-T overlap, meaning that 293 of the genes whose transcription peaks in the LOC-phase show little change in their RNASeq levels, and may represent particularly stable transcripts.

We divided the transformed RNASeq data into the 3 NETSeq clusters and compared the NETSeq and RNASeq metagenes over time (Figure 1C). The largest cluster, (NC-T) containing over 70 % of all genes (3926/5579), strongly resembled the overall profile of transcript expression through the cycle, remaining largely constitutive but peaking slightly in late HOC as cells begin to divide, perhaps reflecting a transient transcript stabilization before homeostasis to gene dosage is restored (Voichek et al., 2016). There are also differences between transcripts and transcription in the oscillating clusters, again suggesting some degree of differential transcript stability. For example, the mean RNASeq range is 4.2-fold in the LOC cluster compared to a mean NET-Seq range of 2.7-fold. For the HOC cluster the mean NET-Seq range is 6.5-fold, compared to a mean RNASeq range of 10.8-fold. Furthermore, in the HOC-T cluster, levels of nascent transcripts return to baseline long before steady-state transcripts do, suggesting that the cytoplasmic transcripts are subject to active degradation (Figure 1C). Interestingly, only the HOC-T cluster can be further subclustered when using the transformed RNASeq data (Figure EV2F), revealing more subtle post-transcription control of transcripts encoding ribosomal proteins (RPGs) or ribosome biogenesis factors (RiBis) (Figure EV2G).

### The majority of the detectable proteome does not cycle

The peaks in levels of steady-state transcripts encoding components for assembly of ribosomes (Figures 2F, EV2G, 3G,H), or enzymes of metabolic pathways (Figure EV3), are widely assumed to drive changes in protein levels which in turn *cause* the changes that characterize this rhythm. For example, levels of ethanol peak during the HOC phase (Figure EV3C, EV Table 1) and levels of transcripts encoding alcohol dehydrogenases also cycle (Figure EV3B). Current models would support a commitment increase in the levels of the alcohol dehydrogenase enzymes to metabolize this ethanol as a carbon source. Similarly, in the HOC phase, when levels of RPG and RiBi transcripts increase (Figure 2F, EV2G), this is interpreted as meaning there is increased production of ribosomes to support protein synthesis required for cell growth and division (Kuang et al., 2014; Murray et al., 2007; Tu et al., 2005; Tu and McKnight, 2006).

**Figure 2.**
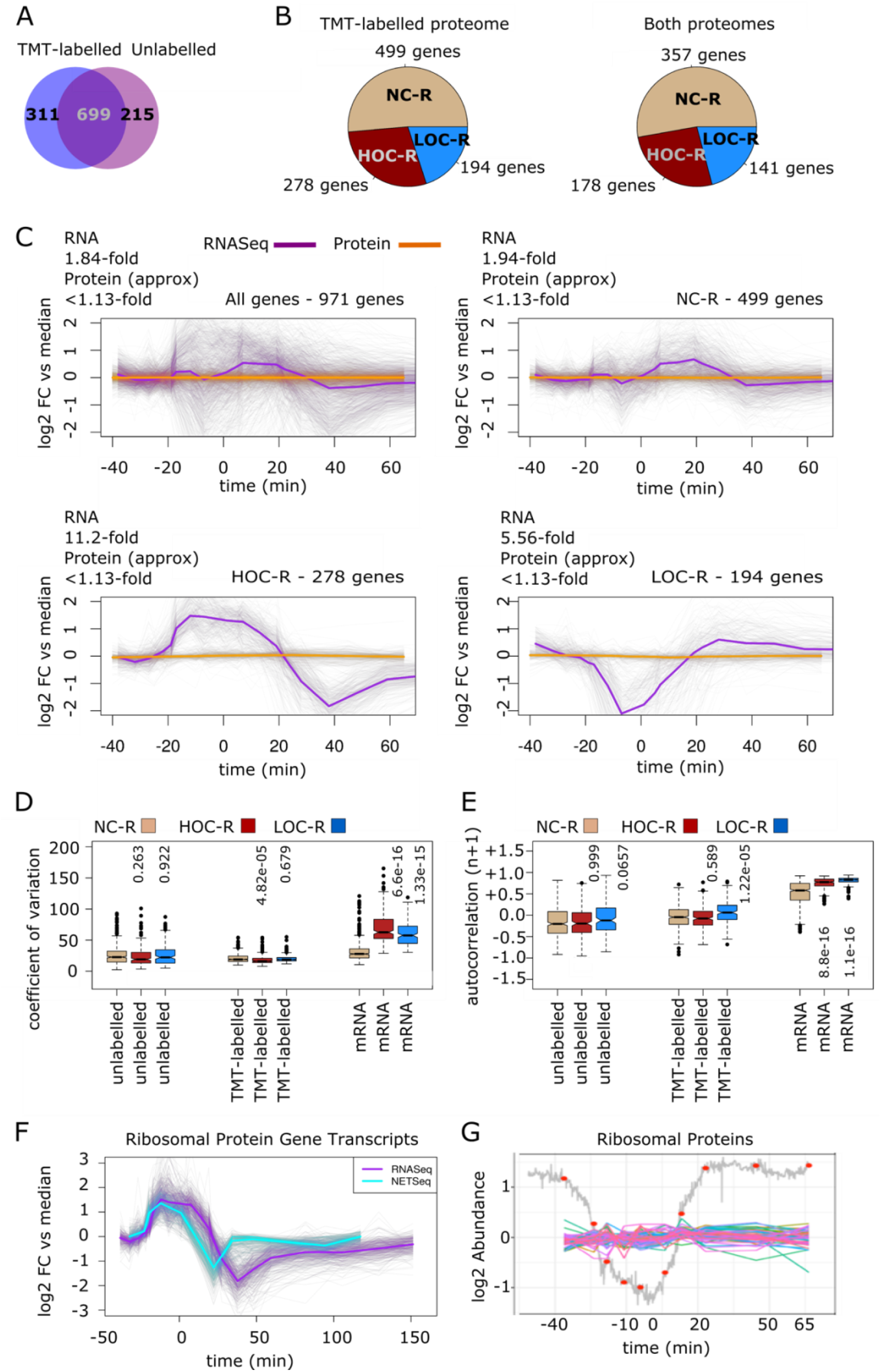
The detectable proteome of the Yeast Metabolic Cycle is static. Two proteomes were generated using TMT-labelled (10 samples) and unlabelled (6 samples) mass-spectrometry from soluble protein extracts from the yeast metabolic cycle. Sampling times (red dots) for the TMT-labelled samples are shown in (**G**). (**A**) A Venn diagram was generated to show the overlap of proteins with coverage over all timepoints between datasets. (**B**) The distributions of the three RNASeq clusters in both proteome datasets were plotted as pie charts. (**C**) A log2 fold-change vs median transformation was applied to the RNASeq data and the TMT-labelled proteomics and both were divided into the 3 RNASeq clusters (R) and plotted over time. (**D**) For each gene, the coefficient of variation of log_2_ fold-change vs median transformed data was calculated for the RNASeq dataset and both proteomes. These values were used to generate box plots splitting the data by dataset and RNASeq cluster. The statistical significance of the difference between the means of NCR genes and HOC or LOC genes was calculated for each dataset using Dunnett’s Test. Raw p-values are displayed. (**E**) The autocorrelation of log2 fold-change vs median transformed values were calculated for each gene for the RNASeq dataset and both proteomes. They were then used to generate box plots splitting the data by dataset and RNASeq cluster. The statistical significance of the difference between the means of NC-R genes and HOC or LOC genes was calculated for each dataset using Dunnett’s Test. Raw p-values are displayed. (**F**) NETSeq and RNASeq read counts for ribosomal protein gene transcripts were LFCM and plotted against time. Levels peak in the HOC phase. TMT-labelled proteome for ribosomal proteins shown as log_2_ abundance over time. The change in the dissolved oxygen (grey line) and exact sampling times (red dots) are shown on the same figure. Sampling times are shown relative to the point of maximum oxygen consumption (0).

**Figure 3.**
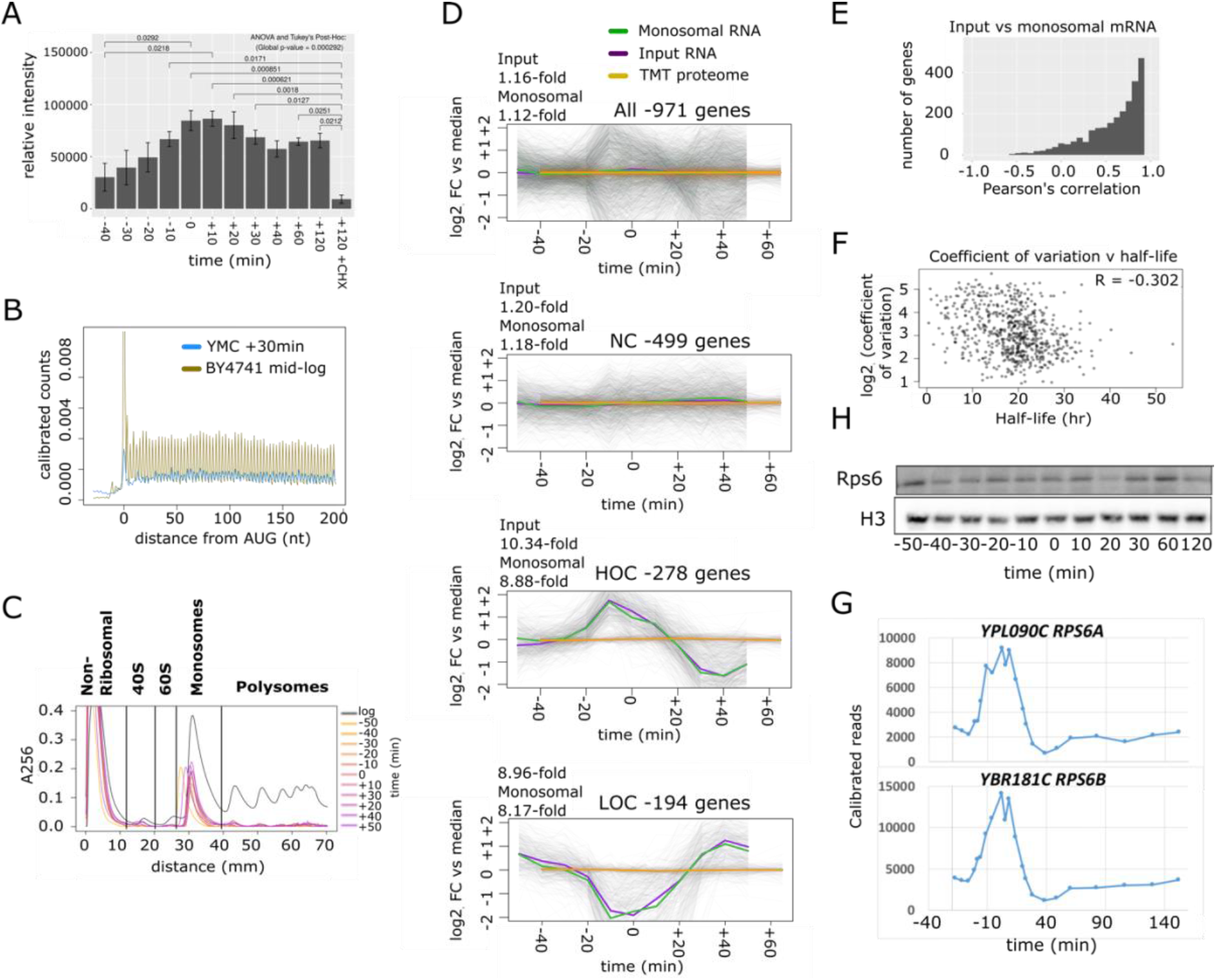
Translation cycles during the YMC. **(A)** Levels of protein synthesis during the YMC, assessed by the incorporation of puromycin and detection using an anti-puromycin antibody. At +120 mins relative to the highest point of oxygen consumption cells were treated with or without cycloheximide to inhibit protein synthesis to control for background levels. n=3, see EV4B,C. Differences were assessed by ANOVA and Tukey’s post hoc test. (**B**) Metagene showing ribosomal occupancy over the first 200nt of transcripts in the YMC sampled at +30 mins relative to the point of highest oxygen consumption and in mid-log cells, see EV4D,E. n=1 (**C**) Polysome profiles from 2 mg of cell lysate at the times shown in the YMC relative to the point of highest oxygen consumption and from a culture in exponential growth (black line), see EV4F,G. n=2 **(D)** Analysis of transcripts associated with monosomes or total RNA. Read counts from RNASeq were LFCM transformed and divided into the 3 clusters and plotted against the TMT proteome from the same transcripts over time where 0 is the time of maximum oxygen consumption n=1 for total data set (**E**) Histogram of the similarity between the input and the monosomal transcripts calculated as Pearson correlation coefficients. (**F**) Plot showing a small negative correlation between transcript half-life (Lahtvee *et al.,* 2017) and the coefficient of variation in transcript levels during the YMC. (**G**) Western blot showing levels of Rps6 protein, with histone H3 as a loading control (representative images for n=5). These samples taken from cycle B in Figure 5A. (**H**) Calibrated normalized read counts for *RPS6A* and *RPS4B* transcripts during the YMC.

To question whether levels of proteins cycle in phase with the transcripts that encode them, we generated two proteomes from synchronized wild-type cells, one with, and one without, Tandem Mass Tag (TMT)-labelling. We used two different methods to ameliorate any potential technical issues with using just one method to assess the proteome. We observed a strong overlap in the representation of the proteome in the two datasets (Figure 2A) and strong representation of all three mRNA clusters (Figure 2B). The TMT-labelled proteome, along with its corresponding transcriptome data, was plotted as LFCM for each RNASeq (R) cluster (Figures 2C, EV3, EV3 appendix 1). Despite the cluster members showing their characteristic profiles at the RNA level, their protein levels remained constant over time (mean range of protein levels, <1.15-fold). This is also shown in the differences between the coefficients of variation (CoV) for mRNA levels and proteins levels of the HOC-R, LOC-R and NC-R clusters (Figure 2D). Furthermore, the autocorrelation of the mRNA was generally strongly positive for all clusters, demonstrating smooth and continuous changes in expression over time, whereas the coefficients for the proteomics data centre around zero, implying that the small changes in the proteomics data are largely the result of statistical noise (Figure 2E). We used Western blots to confirm the proteomics data and show that levels of selected protein, such as tubulin, glyceraldehyde 3 phosphate dehydrogenase, neutral trehalase, histone H3 (Figure EV3 appendix 1) and ribosomal protein S6 (Figure 3H), remain level throughout the rhythm, although in each case, their transcripts levels cycle. Furthermore, the proteomics analysis reveals that levels of ribosomal proteins (Figure 2G) or alcohol dehydrogenases (Figure EV3A) do not change over the cycle, but their transcripts do (Figure 2F; EV3B), suggesting that the models in which changes in protein levels drive the physiological changes during the rhythm are called into question.

### Protein synthesis cycles

We considered three scenarios to explain how the disconnect between the generally invariant proteome and cycling transcriptome arises; selective translation of a subset of the cycling mRNAs, selective protein degradation, or low levels of cycling protein synthesis contributing to a large stable pool of protein (Figure EV4A). Our data support the third model. Firstly, active protein synthesis was detected over most of the rhythm using the puromycin incorporation assay, although it increases over the HOC phase and into the early LOC phase (Figure 3A and EV4B,C). Additionally, calibrated RiboSeq (Ingolia et al., 2009) metagenes revealed that triplet periodic reads were present in RiboSeq libraries from synchronized cells when active protein synthesis is detected (t=+30, where t=0 is the peak oxygen consumption) (Figure 3B, EV4D), supporting active translation across mRNAs (Figure EV4E), albeit at far lower levels than in log-phase (Figure 3B, EV4D). Consistently, ribosome occupancy, assessed using polysome profiling, was dramatically different from exponential growth, primarily occurring as monosomes (Figure 3C and EV4F,G). By sequencing poly(A)-primed RNASeq libraries, prepared from the input and monosomal fractions of the polysome profiles, we showed that translation of genes across the cycle is a simple function of their transcript levels (Figure 3D,E).

To maintain a stable proteome while producing low level rhythmic pulses of new proteins, there must be slow protein turnover rates. In support of this, the median protein half-life at the growth-rates used in our experiments is around 17.2 hours (Lahtvee et al., 2017) and protein half-life negatively correlates with the degree of variation in protein levels through the cycle (Figure 3F). This supports proteins being stable under cycling conditions as the most likely explanation for the invariant proteome (see Figure EV3 and appendices for examples).

As testament to this, the Rps6 protein (Figure 3H) and other ribosomal proteins (Figure 2G) remain constant throughout the cycle, although levels of their steady-state transcripts levels cycle (Figure 2F, 3G, EV2G), peaking in the early HOC phase. The current models whereby increased levels of ribosomal proteins in the HOC phase leads to the synthesis of more protein during the growth phase of the YMC cannot be correct. Thus we sought alternative explanations.

### Post-translational modifications to proteins cycle

Like many biological rhythms, the yeast rhythm alternates between low and high energy states which are coupled to cycling levels of key metabolites such as acetyl CoA and [ATP/ADP] ratios (Cai et al., 2011; Machne and Murray, 2012; Tu et al., 2007). Current models suggest that high ATP levels (das Neves et al., 2010) and histone lysine acetylation at promoters of growth related genes (Cai et al., 2011) drive transcription, which in turn produces a distinct proteome, to determine the different physiological states of the rhythm (Machne and Murray, 2012; Murray et al., 2007; Tu et al., 2005). However, we show here that the detected proteome does not cycle.

An alternative explanation is that protein levels remain constant but post-translational modifications (PTMs) to proteins vary with the changing levels of metabolites during the rhythm to coordinate cellular processes with energy state. Indeed, many metabolites cycle and are often co-factors for enzymes that deposit PTMs. To examine whether PTMs cycle, we focused on phosphorylation and acetylation of selected proteins (Murray et al., 2007; Tu et al., 2007). In the YMC, Rps6 is a stable ribosomal protein which is translated from cycling transcripts (Figure 3H,G). Rps6 is subject to phosphorylation by the nutrient-sensing TORC1/2-signalling pathway and is a marker of ribosome biogenesis (Gonzalez et al., 2015; Yerlikaya et al., 2016). In order to determine whether the levels of phospho-Rps6, and by extension 40S-subunit assembly, vary during the rhythm, we probed Western blots for phospho-Rps6p (S232/S233) (Figure 4A,B, EV5A,B, EV5 appendix 1). Phospho-Rps6 showed a significant enrichment in HOC-phase versus LOC-phase, increasing in abundance several fold. This suggests that the functional phosphorylated form of Rps6 is used in ribosome biogenesis cycles in the YMC. Next we examined whether ribosome assembly cycles during the YMC by monitoring the production of rRNAs, which are also TORC1/2-directed (Honma et al., 2006; Li et al., 2006; Mayer and Grummt, 2006). We used RNA-fluorescence in situ hybridization (FISH) with probes targeted to the transcribed spacer regions allowing nascent rRNA transcript production to be monitored. rRNA transcription rates were low during the LOC-phase and peaked during the HOC, and in addition were expressed in a higher proportion of the cells in the population in the HOC phase (Figure 4C, EV5C, EV5 appendix 2). This suggests that following its phosphorylation, Rps6 plays its role in processing the polycistronic pre-ribosomal RNA (35S) into mature rRNA molecules, facilitating ribosome assembly. Intriguingly, as ribosomal protein (RP) levels were constant over time, we infer the existence of an excess pool of chaperone-bound RPs that are drawn-upon to assemble ribosomes during the HOC-phase. It appears that TORC1/2-signalling and 40S assembly vary across the YMC, explaining the increase in global rates of protein synthesis (Figure 3A) but *NOT as* a result of cycling levels of RPGs and RiBis as is widely assumed in current models. These assumptions arise from the dramatic changes in levels of transcript encoding these proteins.

**Figure 4.**
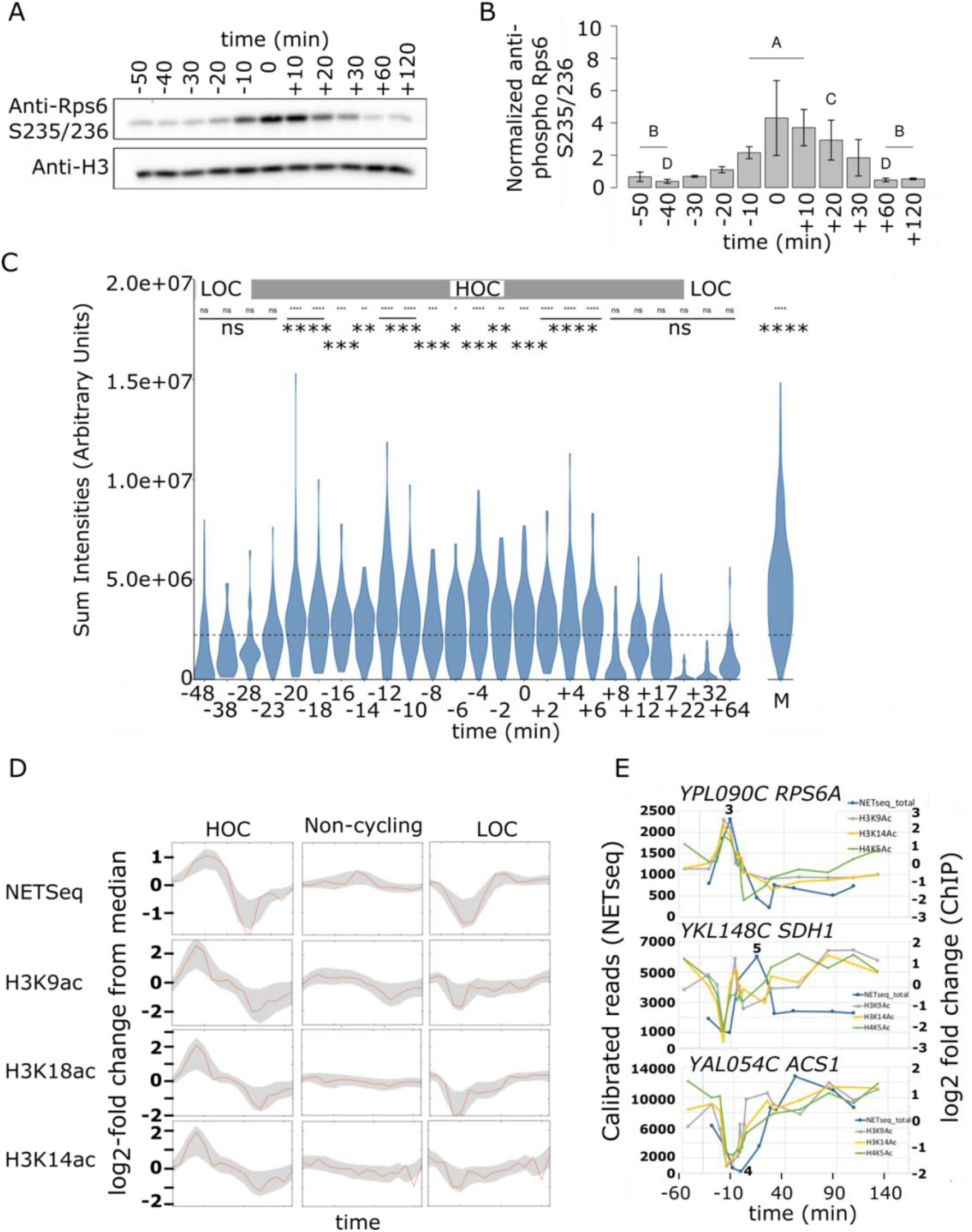
Cycling post-transcriptional modifications to histones and Rps6 influence rhythmic ribosome assembly and transcription the YMC. **(A)** Western blots showing the levels of doubly-phosphorylated Rps6 at S235/S236 and histone H3, as a loading control, at 11 time-points through the YMC. (**B**) Normalisation of Rps6 S235/S236 phosphorylation to histone H3 levels. Group A is significantly greater than group B. Group C is significantly greater than group D using ANOVA and Tukey’s post-hoc test; n=4 (see Figure EV5 appendix 1 for repeats). (**C**) Violin plots showing the sum of the rRNA intensities determined by RNA FISH to spacer regions, allowing assessment of rates of rRNA transcription before processing. Levels of rRNA production peak in the HOC phase of the cycle. Levels are substantially lower than in an exponentially growing mid log culture (M). The sampling time points are shown in minutes relative to the point of lowest dO_2_% in the YMC (0). LOC phase (light grey), HOC phase (dark grey). The dashed line represents the base mean of all the time points sum intensities, excluding M (mid log). Asterisks indicate time points for which rRNA sum intensities are significantly greater than the base mean (p-value<0.05). p-values were calculated by the Wilcoxon test. For Figure EV5 appendix 2 for sampling times, distribution of signal in individual cells and data for 2^nd^ and 3^rd^ repeats. n=3 biological repeats from different chemostat runs, each analysed separately (**D**) Log_2_-fold change from median for levels of H3K9ac, H3K14ac and H3K18ac (by ChIP) at genes whose nascent transcription (NET-Seq) peaks in the HOC or LOC phases of the YMC and the non-cycling gene cluster shown in Figure 1. (**E**) Profiles showing RNAPII density (NETSeq calibrated reads) and levels of H3K9ac, H3K14ac and H4K5ac (log_2_ fold change) at *RPS6A, SDH1* and *ACS1* over the YMC. The point of lowest dissolved O_2_ is 6 mins after sample point 4 (see Figure EV5 appendix 3 for more examples and EV Table 1 for details).

Next we asked why, if the proteome does not cycle, do we observe rhythmic production of some nascent transcripts? Interestingly, in addition to regulating Rps6 phosphorylation and rRNA transcription, TORC1 also regulates dynamic growth-related acetylation of histone H3 and H4 N-terminal lysine residues at promoters, including at ribosomal protein genes (RPGs). Signalling via TORC1 is proposed to rapidly alter chromatin compaction and the potential for transcription by controlling the activity of lysine acetyl transferase, deacetylases and chromatin remodelling ATPase complexes (Damelin et al., 2002; Gowans et al., 2018; Laribee, 2018; Rohde et al., 2004; Rohde and Cardenas, 2003; Workman et al., 2016). Acetyl Co-A-dependent acetylation of histones is known to vary over biological rhythms and is associated with active transcription (Alenghat et al., 2008; Cai et al., 2011; Chatterjee et al., 2011; Etchegaray et al., 2003; Feng et al., 2011; Ferreira et al., 2007; Hirayama et al., 2007; Kuang et al., 2014; Shi and Tu, 2015). Plotting the NET-Seq data alongside published acetyl-H3 ChIP-Seq data (Kuang et al., 2014; Sánchez-Gaya et al., 2018) for the genes in the HOC-T and LOC-T clusters revealed that the changes in expression that they exhibit during the HOC-phase follow changes in histone acetylation status at the same promoters (Figure 4D, EV5 appendix 3, EV Table 1). Consistently, there was no change for the noncycling genes.

To explore changes in acetylation in more detail, we followed acetylation patterns at individual genes whose nascent transcripts peak in early HOC *(RPS6A),* late HOC *(SDH1)* or in LOC *(ACS1)* (Figure 4E, EV5 appendix 3, EV Table 1). This revealed three clear profiles in cycling acetylation: (i) A spike of acetylation early in HOC phase followed by a rapid return to base line with transcription peaking and falling coordinately, (ii) a dip in levels of acetylation early in HOC phase that then spikes in late HOC before returning to base line with transcription peaking after the peak of acetylation or (iii) a dip in levels of acetylation early in HOC phase that rapidly returns to base line and is disconnected from transcription. Generally, the dip in acetylation levels is related to a reduction in the production of nascent transcripts for the late HOC genes and the LOC genes. By contrast, at the early and late HOC genes, the spike in acetylation is tightly related to the production of nascent transcripts. This data supports a temporal association between acetylation and the production of nascent transcripts in the HOC phase. The peak levels of nascent transcripts and histone acetylation at the early HOC promoters is matched by the lowest levels in the late HOC and LOC cluster, suggesting competition for limiting transcription resources focused on the dynamic peaks and troughs in levels of acetylation, which reflect the energy state of the cell. Thus, we propose that the changes in transcription at promoters through the YMC are driven by a shift in acetylation from one class of promoters to the other, titrating limiting PIC components between promoter classes.

## Discussion

The assumption that transcription/translation feedback-loops control and define the phases of the biological rhythms is now being called into question (O’Neill and Feeney, 2014; Ray et al., 2020). Instead, biological rhythms appear to be regulated through changes in metabolism, protein activity and the activity of signaling complexes such as TOR, PKA, SAGA, CK1, GSK3 and Tip60 (Cai and Tu, 2011; Causton et al., 2015; Lee and Tu, 2015). Indeed, regulation of protein activity through metabolites and post-translational modifications makes for much more economical control of biological rhythms than bulk remodelling of the cellular proteome. By demonstrating complete decoupling of transcript and protein levels, we highlight how using the transcriptome as a proxy for gene expression can result in misleading conclusions. Thus, when considering the impact of an internal or external perturbation of gene expression on cellular phenotype, it is critical to measure the effect that it has on protein levels and activity. Additionally, our findings highlight the underappreciated importance of protein stability in controlling gene-expression dynamics, as it is at this stage of gene-expression where we propose that the effect of cycling transcription is silenced. We infer that in non-synchronized cells, it is the transcriptome, not the proteome, that is noisy.

A major question that emerges from this work is “If the proteome remains unaffected by the cycling transcriptome, what is the function of transcriptional cycling across biological rhythms?”. One intriguing possibility is that transcription may have additional roles in cellular physiology, aside from RNA synthesis. Transcription and the chromatin environment are deeply intertwined and extensively remodel one another (Kouzarides, 2007; Skalska et al., 2017). Furthermore, both processes are directly controlled by metabolism as certain core metabolic enzymes such as Hxk2p, and kinases such as TOR, can bind to the DNA via transcription factors and regulate promoter activity (Ahuatzi et al., 2007; Tsang et al., 2010; Vega et al., 2016). Perhaps the role of transcription in biological rhythms is to coordinate and regulate signalling pathways by converting the chromatin into a platform upon which signalling complexes can form and interact with one another to influence other aspects of cellular biology?

Alternatively, it is possible to simply dismiss the cycling transcriptome as a spandrel; a process that occurs not because of any inherent adaptive advantage but because it is a relatively harmless consequence of other beneficial properties of cellular physiology. It is crucial for the cellular transcriptional machinery to be responsive to external nutrient signals such as changes in carbon and nitrogen availability in order to allow the cell to adapt to changes in its environment. Accordingly, it is possible that this nutrient-sensitivity is also enough to drive changes in transcription as a result of the changing intracellular metabolite levels that occur during biological oscillations. That the number of genes expressing cycling transcripts increases as nutrient availability increases (Wang et al., 2015) supports this hypothesis. This hypersensitivity, if present, would not be selected against due to its lack of effect on the proteome. As such, it might be allowed to persist to ensure that cells are able to sense and respond to changes when this is required of them?

Finally, this new view of biological rhythms provides an explanation as to why the cycling of transcripts appear to be linked to the biology of the phase of the rhythm in which it is expressed. For the YMC, this applies particularly to the HOC phase, which is highly responsive to intracellular or extracellular signals particularly the availability of carbon source and conditions that promote growth (Cai et al., 2011). This leads to TORC1 and nutrient-dependent signalling (Kunkel et al., 2019) and by promoting both phosphorylation of Rps6 and the acetylation of the promoter chromatin, creates the *perceived* link between transcription and ribosome biogenesis. Our model suggests that this link is not linear, as widely believed, as ribosomal proteins do not cycle, but is a more complex network. Finally, the cycling of LOC transcripts antiphase to the HOC transcripts can be explained by deacetylation of their promoters although how this is achieved is not yet clear.

In summary, we find that the eukaryotic proteome is able to buffer changes in transcript abundances through slow protein-turnover, highlighting the importance of direct measurements of the proteome and protein activity when making inferences about phenotype. Furthermore, cycling of transcription in the absence of cycling proteins implies that additional functions for transcription may exist aside from gene expression.

## Materials and Methods

### Lead Contact and Materials Availability

As Lead Contact, Jane Mellor is responsible for all reagent and resource requests. Please contact Jane Mellor at with requests and inquiries.

### Method Details

#### Genetic manipulation of yeast strains

All strains used in this study are listed in the KEY RESOURCES TABLE. Genetic manipulation of strains was carried out using the homologous recombination method described (Longtine et al., 1998). For gene deletion strains, PCR products were made containing a selection marker with promoter and terminator sequences flanked at both ends by 40bp of sequence homologous to sequences either side of the region to be deleted. For 3xFLAG tagged strains, first a plasmid was created from plasmid pFA6a-GFP(S65T)-His3MX6 (Longtine et al., 1998) with the 3xFLAG sequence amplified from strain YSC001 by PCR inserted in place of GFP. PCR products were made using this template that consisted of a 40bp sequence homologous to the first 40bp upstream of the stop codon of the gene to be tagged followed by the FLAG sequence, His5 selection marker and 40bp of sequence homologous to a region downstream of the gene to be tagged. 3xHA tagged strains were created in a similar manner using the pFA6a-3HA-KanMX6 plasmid (Longtine et al., 1998). Cells to be transformed were grown to log phase, pelleted, re-suspended in 450 μL 0.1 M LiAc/TE and incubated (> 1 hr, 4°C). 100 μL of cell suspension, 10 μL of gel-extracted PCR product, 10 μL calf thymus DNA (Sigma D8661), 700 μL 0.1 M LiAc/TE, 40% PEG were incubated (30 min, 30°C) then heat-shocked (20 min, 42°C). Cells were pelleted (5 min, 7000 rpm), re-suspended in H_2_O and plated onto appropriate selection media. DNA was extracted from transformants, screened by PCR and confirmed by sequencing.

#### Metabolic Cycles

Fermenters were Bioflo320 (2 l vessel, 1.1 l culture volume – New Brunswick) or Minifor (400 ml vessel, 200 ml culture volume – Lambda) as indicated and cultures were grown in either YMC-YE media (pH 3.5 – ammonium sulphate 5 g/l, potassium dihydrogen monophosphate 2 g/l, magnesium sulphate 0.5 g/l, calcium chloride 0.1 g/l, yeast extract 1 g/l (Difco), glucose 10 g/l, sulphuric acid 0.035 %, antifoam-204 0.05 % (Sigma-Aldrich), iron sulphate 20 mg/l, zinc sulphate 10 mg/l, manganese chloride 1 mg/l, copper sulphate 10 mg/l) or YMC-MD media (pH 3.5 – ammonium sulphate 5 g/l, potassium dihydrogen monophosphate 2 g/l, magnesium sulphate 0.5 g/l, calcium chloride 0.1 g/l, glucose 2.5 g/l, sulphuric acid 0.0067 %, antifoam-204 0.05% (Sigma-Aldrich), iron sulphate 20 mg/l, zinc sulphate 10 mg/l, manganese chloride 1 mg/l, copper sulphate 10 mg/l, biotin 2 ug/l, calcium pantothenate 400 ug/l, folic acid 2 ug/l, inositol 2 mg/l, niacin 400 ug/l, p-aminobenzoic acid 200 ug/l, pyridoxine HCl 400 ug/l, riboflavin 200 ug/l, thiamine HCl 400 ug/l) as indicated. Fermenter runs were initiated with the inoculation of 10 ml starter culture that had been grown overnight to saturation at 30°C. BioFlo320 runs were operated at aeration rates of 1-5 l/min and agitation rates of 600-1000 rpm whilst Minifor runs were operated at an aeration rate of 0.15 l/min and agitation rate of 3.0 Hz. All runs were operated at 30°C and maintained at pH 3.5 through the addition of 0.25 M NaOH. Cells were grown to saturation (OD_600_ 16.0: YMC-YE; OD_600_ 4.0: YMC-MD) and starved for a minimum of 6 hours. Following starvation, continuous culture was maintained through the infusion of fresh media at a dilution rate of 0.082 hr_-1_.

### NET-Seq/RiboSeq

#### Yeast culture

For NET-Seq, 100 ml of cycling *CENPK113-7D rpb3-FLAG* was harvested by filtration onto a 0.45 μm pore size, 90 mm diameter nitrocellulose filter paper (Whatman). Cells were scraped off the filter paper with a spatula pre-cooled in liquid nitrogen and flash frozen in liquid nitrogen. Flash-frozen *S. pombe* cells were added at a mass ratio of 1:5 and cells were ground in 6 cycles of 3 min at a 15 Hz shaking frequency in a 50 mL grinding jar (Retch) with a 25 mm stainless steel ball using a Retsch MM400 mixer mill. The grinding chamber was cooled in liquid nitrogen in between cycles. 1 g of yeast grindate was stored at −80°C. For RiboSeq, 50 ml of cycling *CENPK113-7D rpb3-FLAG* was harvested and ground as above, and 0.5 g of yeast grindate was stored at −80°C.

#### RNase-Protection and Monosome Isolation for RiboSeq

0.5 g of yeast grindate was resuspended in 2.5 ml RiboSeq lysis buffer (20 mM Tris HCl pH 8.0, 150 mM NaCl, 5 mM MgCl_2_, 1% Triton-X100, 100 ug/ml cycloheximide, 1 mM DTT). 250 uL of lysate was digested with 7.5 uL RNasel (Ambion) for 45 min on a nutator (25oC) and monosomes were isolated from the void volume of Illustra MicroSpin S-400 columns (GE Healthcare). Small RNAs were extracted using the miRNeasy kit (Qiagen) and RNA fragments of 26-35 nts in length were isolated using TBU-PAGE (see size-selection).

#### Immunoprecipitation (IP - NET-Seq)

1 g of yeast grindate was resuspended in Lysis buffer A (20 mM HEPES (pH 7.4), 110 mM KOAc, 0.5% Triton X-100, 0.1% Tween 20, 10 mM MnCl_2_, 1x proteinase inhibitors (complete, EDTA-free (Roche)), 50 U/ml SUPERase.In (Invitrogen)). Resuspended grindate was incubated (4°C, 20 min) with 660 U of DNase I (Promega). Insoluble cell debris was pelleted by spinning (16,000 g, 4°C, 10 min). Supernatants were combined and a 20 μL input sample was taken and combined with 20 μL 2 × SDS loading buffer (80 mM Tris-HCl (pH 6.8), 200 mM DTT, 3.2% SDS, 0.1% Bromophenol Blue, 1.6% glycerol)) and added to 0.5 mL of anti-FLAG M2 affinity agarose beads (Sigma) pre-washed with 2 × 10 mL Lysis buffer A (without SUPERase.In). Beads and supernatant were incubated (4°C, 2.5 h) on a nutator before centrifugation (1,000 g, 4°C, 2 min). A 20 μL unbound sample was taken and combined with 20 μL 2 × SDS loading buffer. Beads were washed four times with 10 mL Wash buffer A (20 mM HEPES (pH 7.4), 110 mM KOAc, 0.5% Triton X-100, 0.1% Tween 20, 1 mM EDTA). Beads were incubated (30 min, 4°C) twice with 300 μL of Elution buffer (Lysis buffer A with 1 mg/mL 3 × FLAG peptide (Sigma)). The elution supernatants were combined (a 20 μL elute sample was taken and combined with 20 μL 2 × SDS loading buffer) and RNAs were extracted using a miRNeasy kit (QIAGEN) according to the manufacturer’s instructions.

#### Western blotting

Protein samples were separated by gel electrophoresis on 7.5% or 10% SDS polyacrylamide gels and transferred to nitrocellulose membranes. Membranes were incubated with 5% BSA in TBST (20 mM Tris-HCl (pH 7.5), 150 mM NaCl, 0.1% Tween-20) for 2 hr, then primary antibody in 2.5% BSA in TBST for 1.5 hr, washed, then incubated with rabbit, mouse or rat HRP-conjugated secondary antibody (Sigma) diluted 1:4000 in 2.5% BSA in TBST for 45 min and washed again. Primary antibodies and their dilutions are detailed in the Key Resources Table. Antibody binding was visualized using chemiluminescence (Pierce) and X-ray film.

### Library generation

#### Linker ligation and RNA fragmentation

Immunoprecipitated RNA (3 μg in 30 μL 10 mM Tris-HCl, pH 7.0) was denatured (2 min, 80°C) and placed on ice. A 5’ adenylated, 3’ dideoxy cytosine-blocker cloning linker 5rApp/CTGTAGGCACCATCAAT/3ddC (Integrated DNA Technologies) was ligated to the 3’ ends of RNAs by first dividing the RNA into three microfuge tubes and then adding 10 μL ligation reaction mix to give final concentrations of 50 ng μL^-1^cloning linker 1, 12% PEG 8000, 1 × T4 RNA ligase 2 (Rnl2) (truncated) ligation buffer, 10 U μL^-1^T4 Rnl2 (truncated) (NEB) and incubating for 3 hr at 37°C. Ligated RNA was incubated (95°C, 35 min) with 20 μL Alkaline fragmentation buffer (100 mM NaCO_3_ (pH 9.2), 2 mM EDTA) was used to fragment linker ligated RNA to a narrow size range to reduce size bias of future steps. Fragmentation was not performed for RiboSeq experiments. RNA was precipitated by incubating (30 min, −20°C) with ice cold 500 μL H_2_O, 60 μL 3 M NaOAc (pH 5.5), 2 μL 15 mg mL-1 GlycoBlue (Ambion) and 0.75 mL isopropanol and then spinning (16,000 g, 4°C, 30 min). Pellets were washed with 0.75 mL 80% ethanol, dried (10 min, room temperature) and resuspended sequentially in the same 10 μL 10 mM Tris-HCl (pH 7.0).

#### Size selection

Ligated and fragmented RNA was mixed with 10 μL 2 × TBE-urea loading dye (89 mM Tris-HCl, 89 mM Boric acid, 2 mM EDTA, 12% Ficoll, 7 M Urea, 2.5 mg/ml Orange G), denatured (2 min, 80°C) and run on a 10 well 10% TBE-urea gel (Biorad) (200 V, 32 min). The gel was stained with SYBRGold (Invitrogen) and the region containing 38-95nt fragments excised. This gel piece was divided between three microfuge tubes and physically disrupted. Each was incubated (70°C, 10 min) in 200 μL H_2_O. The tubes were pooled and gel debris was removed using a Costar-Spin-X column (Corning). RNA was precipitated by adding 60 μL 3 M NaOAc (pH 5.5), 2 μL 15 mg mL-1GlycoBlue and 0.75 mL isopropanol, incubating (30 min, −20°C) and then spinning (16,000 g, 4°C, 30 min). Pellets were washed with 0.75 mL 80% ethanol, dried (10 min, room temperature) and resuspended in 10 μL 10 mM Tris-HCl (pH 7.0).

#### Reverse transcription and circularisation

Reverse transcription (RT) was carried out by adding 3.28 μL 5 × FS buffer, 0.82 μL dNTPs (10 mM each) and 0.5 μL 100 μM RT primer (This is phosphorylated at the 5’ end (5 Phos) and contains two 18 carbon spacer sequences (iSp18)), denaturing (80°C, 2 min) then incubating (48°C, 30 min) with 0.5 μL Superase.In, 0.82 μL 0.1 M DTT, 0.82 μL Superscript III (Invitrogen). 1.8 μL 1 M NaOH was added and RNA was degraded (98°C, 20 min). To neutralize, 1.8 μL 1 M HCl was added. cDNA was mixed with 20 μL 2 × TBE-urea loading dye, denatured (3 min, 95°C), and run (loaded in 2 wells, 20 μL per well) on a 10 well 10% TBE-urea gel (Biorad) (200 V, 50 min). The gel was stained with SYBRGold (Invitrogen) and the regions containing the RT product excised. Both gel pieces were physically disrupted and incubated (70°C, 10 min) in 200 μL H_2_Oin separate tubes. Gel debris was removed and tubes were pooled. cDNA was precipitated by adding 25 μL 3 M NaCl, 2 μL GlycoBlue and 0.75 mL isopropanol, incubating (−20°C, 30 min) and then spinning (16,000 g, 4°C, 30 min). The pellet was washed with 0.75 mL 80% ethanol, dried (10 min, room temperature) and resuspended in 15 μL 10 mM Tris-HCl (pH 8.0).

The RT product was circularized by incubating (60°C, 60 min) with 2 μL 10 × CircLigase buffer, 1 μL 1 mM ATP, 1 μL 50 mM MnCl_2_ 1 μL CircLigase (Epicenter). The enzyme was heat-inactivated at 80°C, 10 min.

#### PCR amplification

16.7 μL 5 × HF Phusion buffer, 1.7 μL dNTPs (10 mM), 0.4 μL 100 μM Barcoded primer (A, B, C, or D), 0.4 μL 100 μM Primer 1, 59.2 μL H_2_O, 0.8 μl Phusion polymerase (NEB) was added to 5 μL circularized DNA. This was divided (16.7 μL per tube) between five 0.2 mL tubes. PCR reactions were heated (98°C, 30 s) and then submitted to 7 temperature cycles (98°C, 10 s; 60°C, 10 s; 72°C, 10 s). At the end of cycle 3 and each subsequent cycle one tube was removed and placed on ice for NETSeq, and adjusted empirically for RiboSeq. 3.4 μL DNA loading dye (1.5 g Ficoll 400, 25 mg Orange G in 10 mL H_2_O) was added to each reaction and run on an 8% TBE gel (Invitrogen) (180 V, 55 min). The gel was stained with SYBRGold and the PCR product excised from the PCR reaction with the highest unsaturated signal without higher molecular weight products. The gel piece was physically disrupted and incubated (room temperature, overnight, with agitation) in 0.67 mL DNA soaking buffer (0.3 M NaCl, 10 mM Tris-HCl pH 8.0, 1 mM EDTA). DNA was precipitated by adding 2 μL GlycoBlue and 0.68 mL isopropanol, incubating (−20°C, 30 min) and spinning (16,000 g, 4°C, 30 min). The pellet was washed with 0.75 mL 80% ethanol, dried (10 min, room temperature) and resuspended in 10 μL 10 mM Tris-HCl (pH 8.0).

#### Sequencing

Samples were submitted to the Harvard Biopolymers facility who determined DNA quality and quantity using the TapeStation (Agilent) and qPCR and then sequenced samples on an Illumina HiSeq 2000 machine or a NextSeq500 machine. 50nt were sequenced from one end using the sequencing primer. Barcoded samples were pooled so that 2-3 samples were multiplexed per lane.

#### Polysome fractionation

Samples were resuspended in 400 uL of polysome lysis buffer (20 mM HEPES pH 8.0, 50 mM KCl, 10 mM MgCl_2_, 1% Triton-X100, 100 ug/ml cycloheximide, 1 mM DTT, complete EDTA-free protease inhibitors – Roche) plus 500 uL acid-washed glass beads (Sigma-Aldrich) and subjected to 8 cycles of lysis in a MagNALyser (Roche – 6000 rpm, 1 min on, 2 min off). Lysate was transferred to fresh microfuge tubes and clarified by centrifugation (13,000 rpm, 10 min, 4°C).

Lysate containing 2 mg of protein was loaded onto the top of a 7-47 % sucrose gradient in polysome gradient buffer (20 mM HEPES pH 8.0, 50 mM KCl, 10 mM MgCl_2_, 100 ug/ml cycloheximide) and macromolecular complexes were separated by velocity gradient ultracentrifugation (SW41Ti – Beckman, 39,000 rpm, 2 hr, 4oC). Fractions were collected using Biocomp Gradient Station and pooled to obtain separate monosomal and polysomal fractions. RNA was extracted from pooled fractions and input samples (miRNeasy – Qiagen) for 3’-end sequencing (QuantSeq for Ion Torrent – Lexogen). Fixed quantities of *S. pombe* RNA were added prior to RNA extraction for the purposes of internal calibration (polysomal – 1 ug_*pombe RNA*_/mg_*Iysate*_, monosomal – 1 ug_*pombe RNA*_/mg_*Iysate*_, input – 30 ug_*pombe RNA*_/mg_*Iysate*_). No polysomal libraries were generated in this study.

#### 3’-end Sequencing of Total mRNA

1 ml of cycling *CENPK113-7D rpb3-FLAG* in YMC-YE media (OD_600_ approx. 16.9) was combined with 2.8 OD_600_ of *S. pombe* and total RNA was isolated by hot acid-phenol extraction. 500 ng of total RNA was used to generate 3’end Seq libraries using the QuantSeq for Ion Torrent kit (Lexogen) and libraries were sequenced on the Ion Proton platform.

#### Extraction of Total Protein

1 ml of cycling *CENPK113-7D rpb3-FLAG* in YMC-YE media (OD_600_ approx. 16.9) was centrifuged (7,000 rpm, 1 min), the supernatant was removed and the pellet was flash-frozen in N2(l).Pellets were thawed, washed twice in 1x PBS (5000 rpm, 3 min, 4oC) and resuspended in proteome lysis buffer (50 mM Tris HCl pH 8.0, 8 M urea, 75 mM NaCl, 100 mM sodium butyrate, cOmplete protease inhibitors – Roche). Cells were then mechanically lysed (MagNAlyzer Roche – 6000 rpm, 1 min on, 2 min off) in the presence of 300 uL acid-washed glass beads (Sigma-Aldrich).

#### Unlabelled Proteome

Protein samples were digested according to the FASP procedure described in Wisniewski *et al.* 2009 (Wiśniewski et al., 2009). After digestion, peptides were separated by nano-flow reversed-phase liquid chromatography coupled to a Q Exactive Hybrid Quadrupole-Orbitrap mass spectrometer (Thermo Fisher Scientific) using HCD fragmentation. In brief, peptides were loaded on a C18 PepMap100 pre-column (300 μm i.d. x 5 mm, 100Å, Thermo Fisher Scientific) at a flow rate of 12 μL/min in 100% buffer A (0.1% FA in water). Peptides were then transferred to a 50cm in-house packed analytical column heated at 45°C (75 μm i.d. packed with ReproSil-Pur 120 C18-AQ, 1.9 μm, 120 Å, Dr.Maisch GmbH) and separated using a 60 min gradient from 15 to 35% buffer B (0.1% FA in ACN) at a flow rate of 200 nL/min. Q Exactive survey scans were acquired at 70,000 resolution to a scan range from 350 to 1500 m/z, AGC target 3e6, maximum injection time 50 ms. The mass spectrometer was operated in a data-dependent mode to automatically switch between MS and MS/MS. The 10 most intense precursor ions were submitted to HCD fragmentation using an MS/MS resolution set to 17 500, a precursor AGC target set to 5e4, a precursor isolation width set to 1.5 Da, and a maximum injection time set to 120 ms. MS data were searched against the *Saccharomyces cerevisiae* UniProt Reference database (retrieved 13/01/17) using MaxQuant, version 1.5.0.35 (Tyanova et al., 2016); precursor mass tolerance was set to 20 ppm and MS/MS tolerance to 0.05 Da. Enzyme specificity was set to trypsin with a maximum of two missed cleavages. False discovery rate for protein and peptide spectral matches was set at 0.01. Acetylation (Protein N-term) and oxidation (M) were set as variable modifications and carbamidomethylation (C) as a static modification.

#### Tandem Mass Tag (TMT)-labelled proteome

Samples were reduced with DTT (final concentration 5 mM), alkylated using Iodoacetamide (final concentration 20 mM) for 30 min each and methanol/chloroform precipitated. In brief, the sample solution (200uL) was mixed with 600 uL methanol, 150 uL chloroform and 450 uL ddH2O. After 3 min centrifugation at 17000xg the upper phase was taken off carefully and another 450 uL methanol were added. After another centrifugation step for 5 min supernatants were taken off completely and discarded. The protein pellets were resuspended in 300 uL 200 mM HEPES, pH 8.5 and digested with 1:100 trypsin over night at 37 C.

Peptide concentrations were measured using Pierce colorimetric peptide assay and the samples were normalized to 225ug of peptides in 300 uL 200 mM HEPES, pH 8.5. 225 ug peptides were labelled with 0.8 mg of a TMT10plex label for 1h and then quenched with 8 uL 5% Hydroxylamine for 15 min. All ten samples were combined, desalted on SepPak Plus cartridges (Waters) and dried down in a vacuum centrifuge.

Peptides were separated on an EASY spray column (ES803, Thermo Fisher Scientific) and analysed on a Dionex Ultimate 3000/Orbitrap Fusion Lumos platform (both Thermo) as described earlier (Davis et al., 2017). We used the MultiNotch MS3 method (McAlister et al., 2014) for generating spectra for identification and quantitation of peptides or phosphopeptides.

MS data were analysed in Proteome Discoverer 2.1. Proteins were identified with Sequest HT against the *Saccharomyces cerevisiae* UniProt Reference database (retrieved December 2014). Mass tolerances were set to 10 ppm for precursor and 0.5 Da fragment mass tolerance. TMT10plex (N-term, K), oxidation (M) and deamidation (NQ) were set as variable modifications and carbamidomethylation (C) as a static modification.

#### MS Data availability

Data are available via ProteomeXchange with identifier PXD019669.

#### Ethanol Assay

Levels of ethanol in the fermenter during the YMC were monitored using the kit from Megazyme (catlog number: K-ETOH).

#### Puromycylation Assay

>3 x 10^8^ cells were permeablised in 0.0003% SDS, then 0.4 mg/ml (final concentration 750μM in H_2_O) puromycin was added and cells were incubated for 4.45 mins at 30°C with shaking. Cells were spun for 15s, supernatant was removed and pellets were frozen before protein extracts made.

#### Western Blotting

1 ml samples from a cycling culture were centrifuged (7000 rpm, 15s), the supernatant was removed and the pellets were flash-frozen in N_2(l)_. Pellets were thawed, resuspended in 600 uL loading buffer (50 mM Tris.HCl pH 8.0, 4 M urea, 100 mM DTT, 10 % glycerol, 2 % SDS, 0.05 % bromophenol blue) and mechanically homogenized using acid-washed glass beads for 3 min (Sigma-Aldrich) then incubated at 95°C for 5 min. Proteins were resolved by SDS-PAGE and transferred to PVDF membranes by wet-transfer blotting (25 V, 16 hr, 4°C). Membranes were blocked for 1 hour in TBST + 5 % BSA, before being treated with primary antibodies in TBST + 2.5 % BSA under the indicated conditions then 3x washed with TSBT. Secondary antibody treatments were performed for 45 min at a dilution of 1:2000 in TBST + 2.5 % BSA at 25°C and followed with 3x washes in TBST. Proteins were detected by chemiluminesence which was recorded either using x-ray film or using an iBright western blot imager (ThermoFisherScientific).

#### RNA fluorescence in situ hybridization (RNA FISH)

Except for the spheroplasting and RNAseA treatment, RNA FISH experiments were performed as described for the ‘endogenous set’ in (Brown et al., 2018). Appropriate volumes of both mid-logarithmic and chemostat cultures (in YMC-MD media, dilution rate of 0.072 h^-1^) were harvested, pelleted and fixed. The amount of cells needed was then pelleted and left on ice to facilitate sampling through an entire cycle of the YMC. Once gathered, pellets were re-suspended in 500μl of spheroplast buffer (Buffer B, 36 μM B-ME) and kept at 30°C 20 min. Cells were then centrifuged, washed (Buffer B, 5 μM B-ME) then spheroplasted using Zymolyase in wash buffer (10mg/ml final concentration, 20.000 U/g, MP Biomedicals) and left at 30°C 4 min, until the majority of the cells were spheroplasted, as observed by microscopy. Integrity of the cells was checked later on, using brightfield microscopy. Cell were washed with buffer B (1.2 M sorbitol, 100 mM K2HPO4, pH 7.5) and attached to coverslips. Samples were rehydrated with 2 ml of 2× SSC for 5 min at room temperature. Coverslips destined to be treated with RNAse A (negative controls) were then transferred to new 12-well tissue culture dishes to avoid contamination. Samples were treated with 2 ml of pre-warmed 2xSSC with or without RNAseA (50 μg/ml final concentration, Merck) and incubated 1h30 min at 37°C. ribosomal RNA probes labelled with Quasar 570 were used at a final concentration of 10nM in hybridization buffer per coverslips. Concerning microscopy, 21 0.2 μm z stacks were imaged with an exposure time of 0.035 s (50 %T) and 0.05 s for TRITC and DAPI channels, respectively. RNA-FISH experiments were repeated 3 times with independent biological samples.

##### rRNA FISH microscopy images

Brightness and contrast of all fluorescence images (examples shown in Figure EV5C) were adjusted to the same levels using FIJI (Schindelin et al., 2012). Background was removed by subtracting a “reference image” of the field taken without any cells from the bright-field images.

##### rRNA FISH probes design

46 DNA probes of ~20 nt, labelled with Quasar 570, were designed to bind both external and internal transcribed spacers regions of the 35S rRNA (RDN37-2: SGD). The probes were obtained from Stellaris (Stellaris probe designer https://www.biosearchtech.com/support/tools/design-software/stellaris-probe-designer).

##### rRNA FISH – Quantification and Analysis

RNA-FISH quantification and preliminary analysis was based on the protocol followed in (Brown et al., 2018) and adapted to detect the larger foci that were produced when labelling rRNA; an identical set of software packages were used.

##### Deconvolution and background subtraction

Deconvolution and background subtraction were performed as in (Brown et al., 2018)

##### Automated Nuclei Detection

In the detection of nuclei, the protocol followed here differed most significantly from (Brown et al., 2018). Initial nuclei detection was identical to that for the ‘endogenous set’ (Brown et al., 2018). The nuclei were identified in 3D using the DAPI channel of the images. The DAPI signal of each image was scaled to the minimum and maximum intensities observed in each image, *P_i_* = (*P_i_* – *min*(*p*))/(*max*(*p*) – *min*(*p*)), where *i* runs over all pixels in the image and *min* (*p*) and *max*(*p*) denote the minimum and maximum pixel intensities of the set, respectively. These scaled images then had a Gaussian filter applied (Matlab function imgaussfilt3 with a smoothing kernel standard deviation value of 2.5). Each image was divided using two threshold values (Matlab multithresh) and segmented into three levels (Matlab imquantize). Each segmented image was then restricted to only pixels from z planes 3 to 17, inclusive. New logical images were constructed from the restricted pixels with the value equal to the highest segment value, 3. The z-stacks were cycled through and any holes were filled (Matlab imfill, flag ‘holes’). All 3D-contiguous regions (with standard 26 connectivity) of pixel value 1 with more than 40,000 or fewer than 200 pixels in them were removed by being set to zero value. The remianing regions were separated and labelled (Matlab bwlabeln) to form the 3D nuclei masks (which will later be filtered depending on cell detection). To aid in the cell detection, these masks were used to generate 2D nuclei ‘seeds’ by making a logical matrix from the sum of the nuclei masks in the z-direction and then labelling (Matlab bwlabel with 8 connectivity).

##### Automated Cell Detection

Barring the use of the alternatively-detected nuclei seed data described above, the automated cell detection was performed identically as in (Brown et al., 2018) for the ‘endogenous set’.

##### Foci classification and rRNA quantification

Following inspection of the distributions of rRNA-channel pixel intensities for all collected data sets and comparison with the same conditions but RNaseA treated, a uniform threshold intensity of 1000 was selected for pixels to count towards rRNA foci. Contiguous regions of intensity greater than the threshold were identified and labelled (Matlab bwlabeln with standard 26 connectivity). The following metrics were used to quantify the foci and cells. The number of separate foci in each cell. The volume of each cell in pixels. The mean intensity of all rRNA foci pixels in each cell. The standard deviation of all rRNA foci pixels in each cell. The total (summed) intensity of all rRNA foci pixels in each cell. The maximum pixel intensity of all rRNA foci pixels in each cell. The number of rRNA foci pixels that were contained in the nucleus of each cell. The total (summed) intensity of all rRNA foci pixels that were contained in the nucleus of each cell.

##### rRNA FISH Downstream analysis

Further analysis was performed using R (R Core Team (2019). R: A language and environment for statistical computing. R Foundation for Statistical Computing, Vienna, Austria. URL https://www.R-project.org/.) Violin plots of the total (summed) intensity of all rRNA foci pixels in each cell for samples taken at different time points through the YMC were constructed for each repeat. QQ plots showed that the rRNA sum intensities were not normally distributed. Therefore, Wilcoxon tests were performed on the rRNA sum intensities to assess which sampling time point showed higher rRNA levels than the base mean of all the time points sum intensities, excluding the sample taken in the middle of the logarithmic phase.

### Software and Algorithms

#### Quantification and Statistical

##### Analysis Aligning NGS data

RiboSeq data was aligned using Tophat (https://ccb.jhu.edu/software/tophat/index.shtml) to a combined *S. cerevisiae/S. pombe* genome using the following parameters: -N 1 -I 2000 -g 5 --b2-N 1 --b2-L 28 -G (gtf file) -no-novel-juncs, and uniquely aligned reads were extracted using samtools view -bq 49. Files were converted to .bed and .bedgraph format using bedtools for further analysis.

NET-Seq data was aligned using bowtie (Langmead, 2010) using the following parameters: – q -p 16 -S -m 1 -n 1 -e 70 -l 28 -k 1 --best -phred33-quals, converted to a .bam file using samtools view -Sb, then into .bed and .bedgraph files using bedtools (citation).

3’-end Seq data was aligned to a combined *S. cerevisiae/S. pombe* genome using STAR (Dobin et al., 2013) with the following parameters: --runThreadN 8 --genomeDir [STARfiles] – -readFilesIn [input_file] --outFilterType BySJout --outFilterMultimapNmax 20 -- alignSJoverhangMin 8 --alignSJDBoverhangMin 1 --outFilterMismatchNmax 999 outsFilterMismatchNoverLmax 0.08 --alignIntronMin 13 --alignIntronMax 2482 -- outFileNamePrefix [output_file]. Uniquely aligned reads were extracted using samtools view -Sbq 20 and converted to .bed format using bedtools bamtobed for further analysis. Raw counts tables were generated from .bed files in R using the summarizeoverlaps function.

##### Library Size Calibration

For 3’-end Seq data from total RNA, library sizes were calibrated to *S. pombe* counts as an internal control, using the estimateSizeFactorsforMatrix command from DESeq2. 3’-end Seq from gradient-fractionated RNA was normalized in the same way, but to the *S. cerevisiae* library sizes (i.e. without internal calibration).

For NET-Seq data, library sizes were calibrated to *S. pombe* counts as an internal control by multiplying by the following normalization factor [(mass of *S. pombe* paste)/(total *S. pombe* reads * mass of *S. cerevisiae* paste)].

For RiboSeq data, library sizes were calibrated to *S. pombe* counts as an internal control.

##### Cluster Analysis

To cluster genes based on their temporal profiles at the transcription, transcript or protein level respectively, calibrated counts for each gene were transformed to obtain log_2_ foldchange in expression over time versus the median. Transformed data was then partitioned using k-means++ clustering into the optimal number of clusters, as determined by the GapStatistic methodologies (https://doi.org/10.1111/1467-9868.00293).

##### Metrics for Dispersion and Oscillation

The coefficient of variation (mean/standard deviation) of signal over time was used to quantify the relative amplitude of cycling variation in potentially cycling 3’-end Seq and proteomics datasets. Signals that oscillate with a large amplitude will have a larger standard deviation over time whereas their mean signal should be invariate. To determine whether variation in signal was continuous or stochastic over time, the autocorrelation at offset +1 was calculated.

##### Western Blot Quantification

Band intensity for Western blotting experiments was quantified using ImageJ and data was analysed in R using ANOVA and Tukey’s post hoc significant difference test. A p-value of less than 0.05 was considered statistically significant.

## Data and Code Availability

All sequencing data generated in this study is available at: https://www.ncbi.nlm.nih.gov/geo/query/acc.cgi?acc=GSE138023.

## KEY RESOURCES TABLE

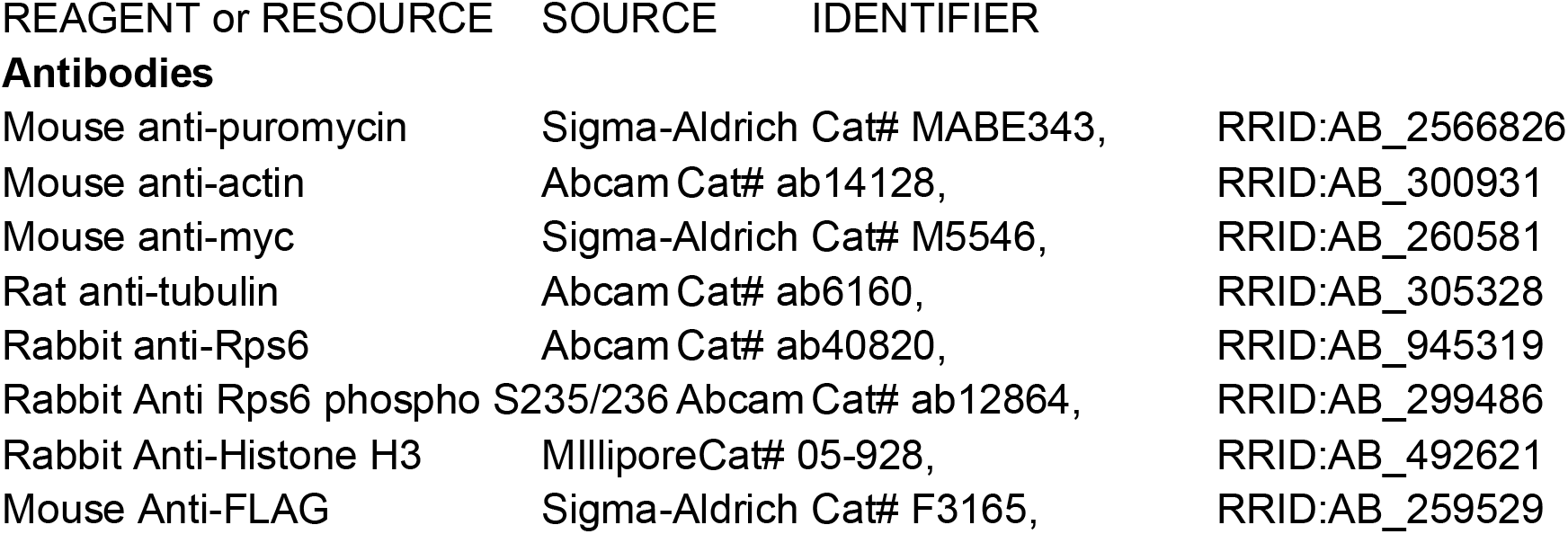
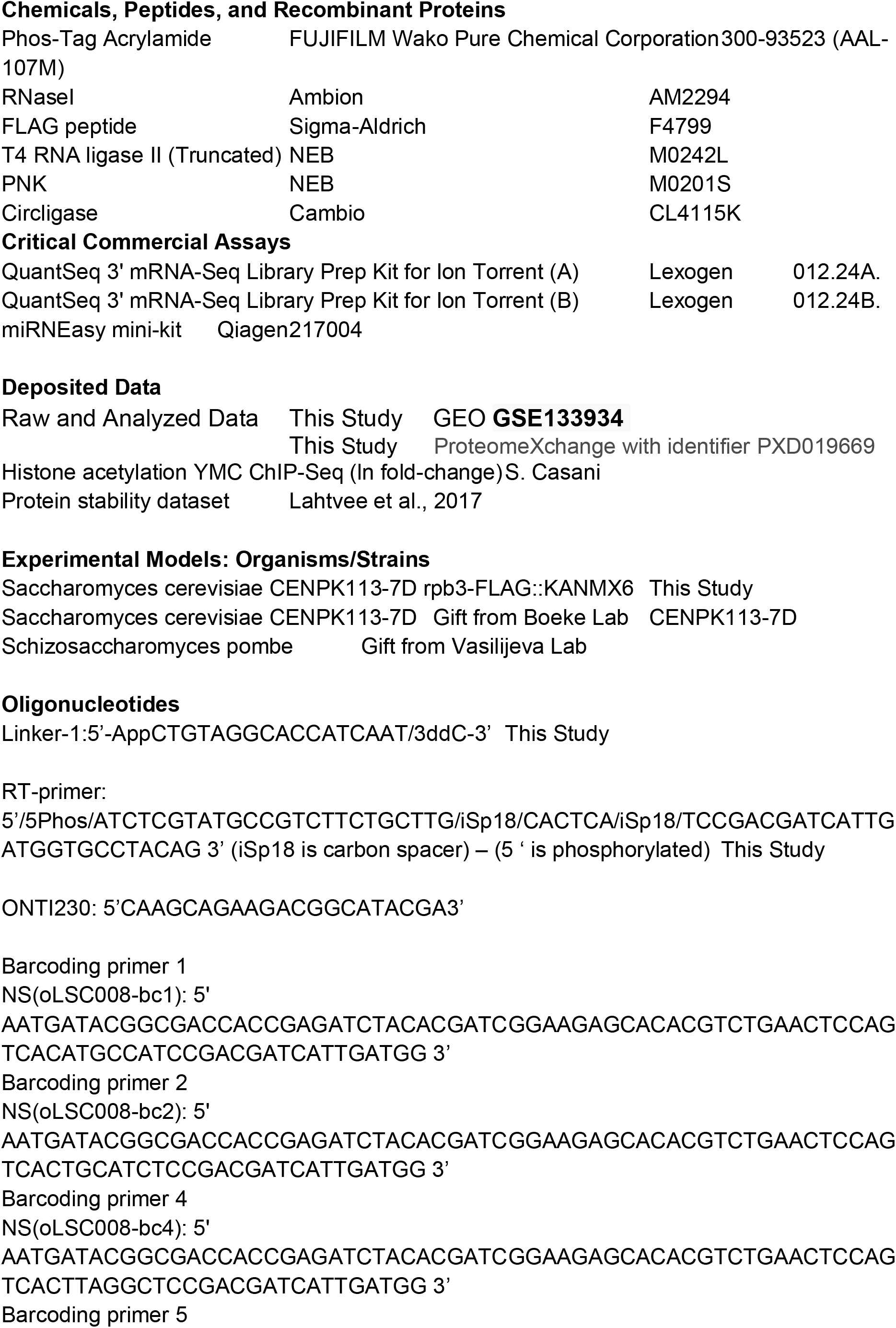
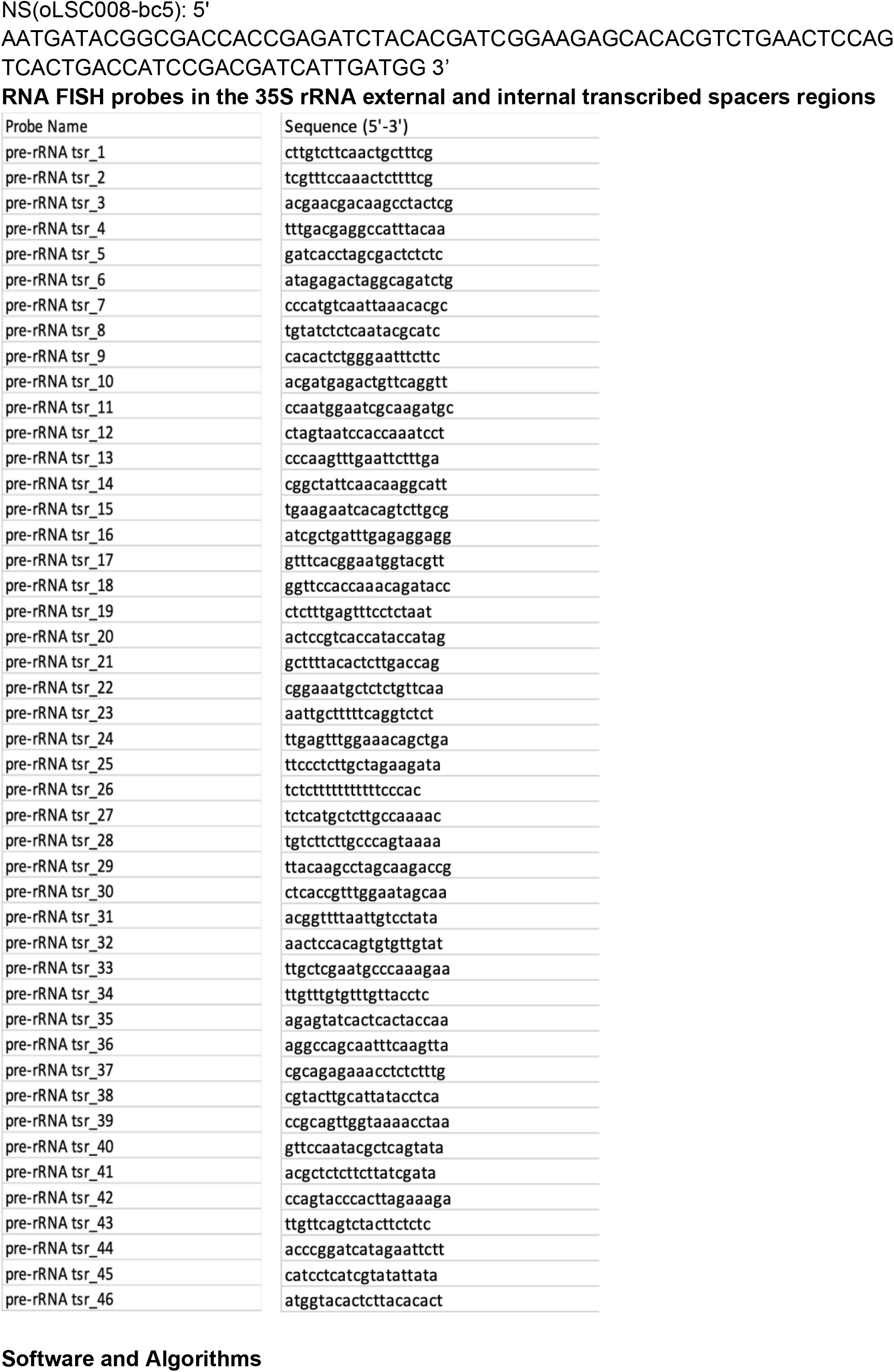
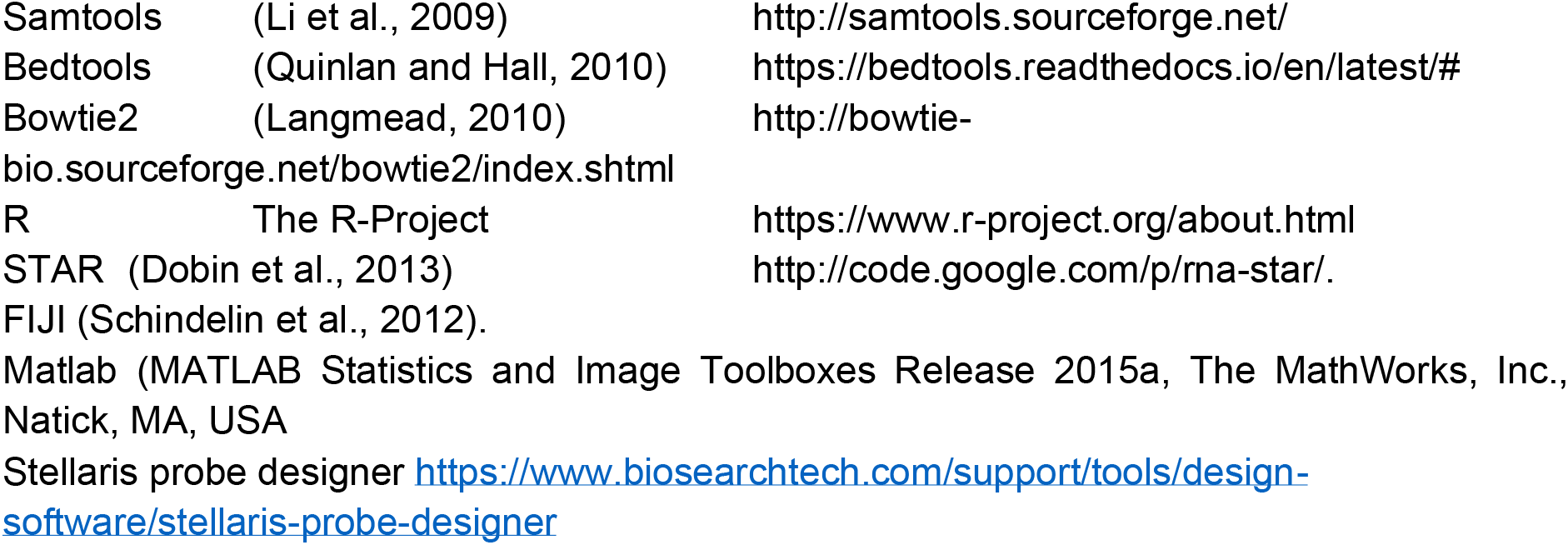

## Acknowledgements

We thank the J.M. lab, the A.A. lab and Andre Furger and his lab for critical discussions; and Ilan Davis and Micron Oxford for microscopy support. This work was funded by the Biotechnology and Biological Sciences Research Council (BB/P00296X/1 and BB/S009035/1 to J.M.); the Leverhulme Trust (RPG-2016-405 to J.M.); a Royal Society University Research Fellowship to AA (UF120327 URF/R/18001); a studentship funded by the European Commission (ITN PEP-NET to MW). We thank the Micron Advanced Bioimaging Unit (supported by Wellcome Strategic Awards 091911/B/10/Z and 107457/Z/15/Z) for their support & assistance in this work.

## Author contributions

JEF ribosome profiles, polysome profiles, RNA-seq, puromycylation, and bioinformatics, data analysis and interpretation, manuscript writing; SX NETSeq, RNASeq, proteomics and data analysis; SCM bioinformatics; MW RNA FISH and downstream analysis; JU-A Rps6 phosphorylation; CG proteomic bioinformatics and analysis; AT, puromycylation and Rps6 phosphorylation; ER puromycylation assay development; RH, TMT-proteomics; BK, TMT-proteomics; SL, label-free proteomics; PC Proteomics analysis; AA RNA FISH scripts and analysis; RF TMT-proteomics; JM conceived project, funded project, data analysis and interpretation, manuscript writing.

---

**Figure EV1.**
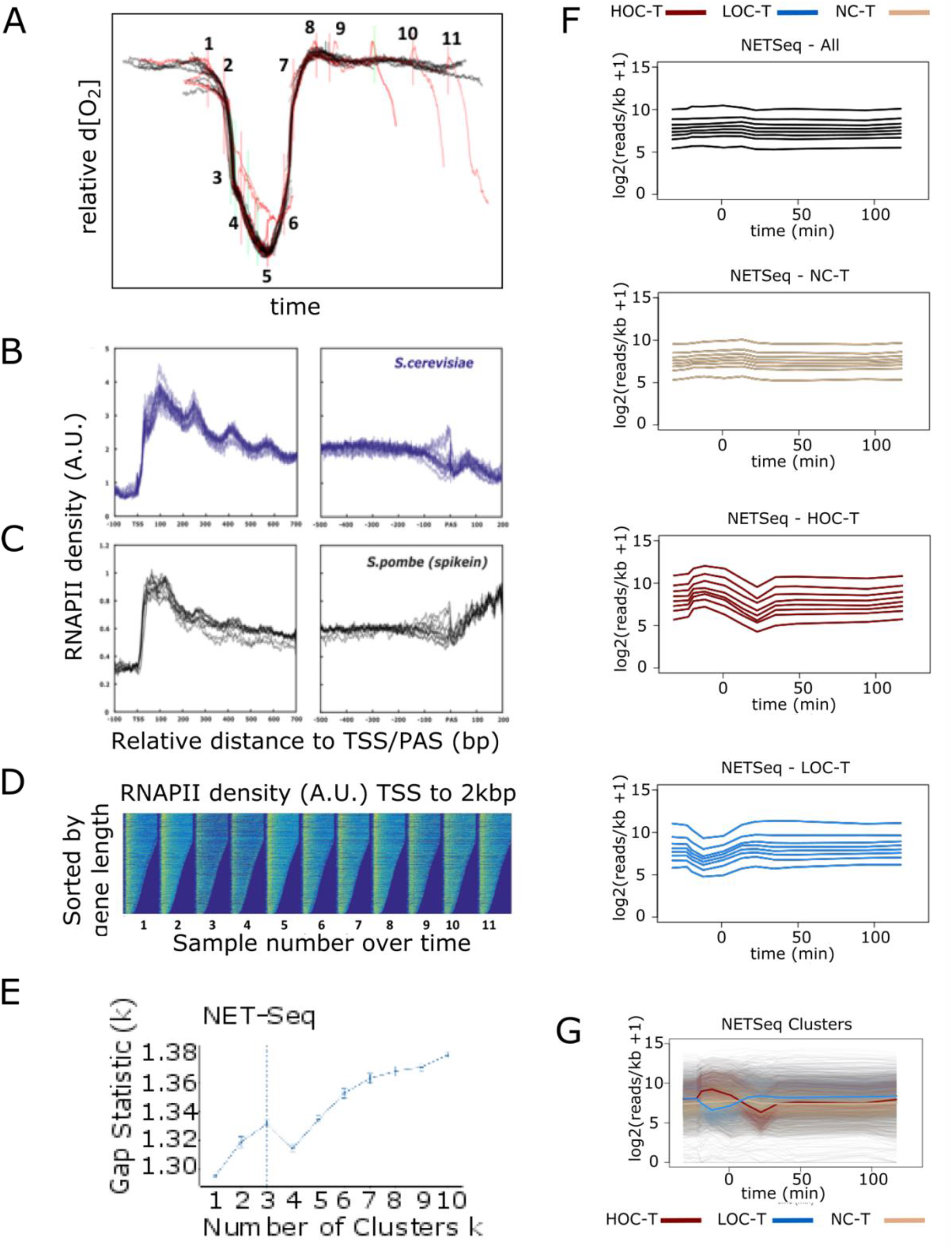
Generation and Analysis of NETSeq data. (**A**) For each time point a 100ml sample from a total volume of 1.1l was taken from a separate cycle. Sampling with large volumes disrupted the cycle causing subsequent entry into an HOC phase meaning that consecutive samples could not be taken in a single cycle. The oxygen consumption profiles for the 11 cycles in which a sample is taken are overlaid. 1ml samples for RNASeq were taken from the same samples to allow direct comparison between the NETSeq data and other RNASeq and ChIPSeq data (Kuang et al., 2014), see Figure 4D and EV5. Detailed sampling times are given in Table 1. The times in all the figures using this data are presented relative to the point of highest oxygen consumption as time 0 to enable direct comparison between different experiments. The reproducibility of the cycle length is evident from the overlays of the 11 cycles. (**B,C**) Proximal and distal metagenes for each 11 NETSeq samples plotted as read counts representing RNAPII density, expressed as arbitrary units for S.cerevisiae YMC samples (**B**) and the S.pombe spike-in samples (**C**). (**D**) Heatmaps showing the *S.pombe* normalized RNAPII density over the first 2kbp of genes from the transcription start site (TSS) sorted by length for each of the 11 NETSeq samples from *S.cerevisiae.* (**E**) Gap statistic analysis of NETSeq data during the YMC indicating 3 is the optimal number of clusters. (**F**) The NETSeq counts tables were normalized to genelength, then log2(x+1) transformed, divided into the NETSeq clusters, then divided into octiles based on median expression level, and plotted over time. Profiles were colour-coded based on cluster membership (red – HOC-T; blue – LOC-T, tan – NC-T). (**G**) Mean profiles for each cluster were also plotted.

**Figure EV2.**
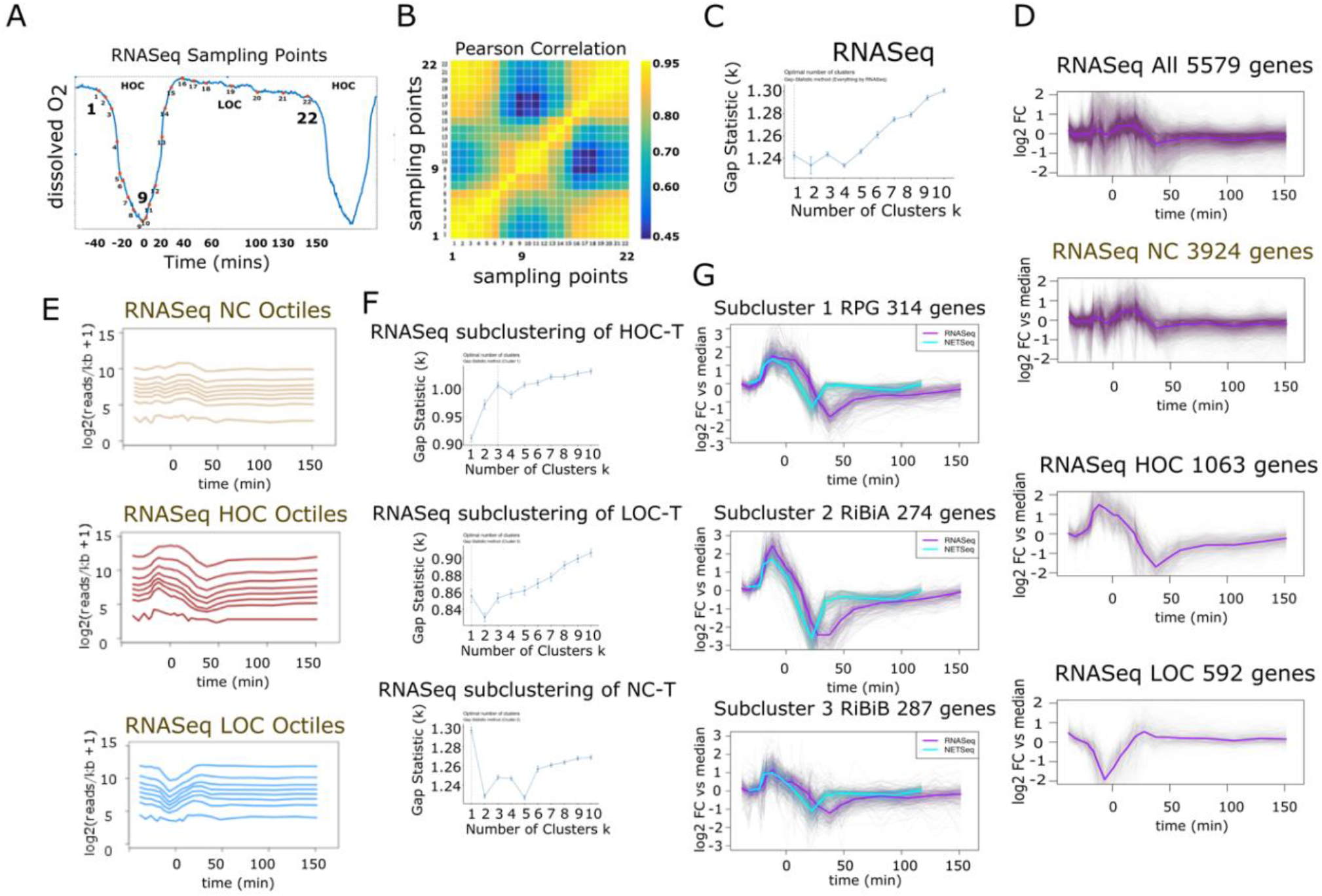
Generation and Analysis of RNASeq data. **(A)** 22 sampling points (red dots) for the RNASeq data shown relative to the oxygen consumption rate obtained from calibrated dO_2_ levels in the culture. 1ml samples for RNASeq were taken during one cycle at the times indicated See EV Table 1 for details. (**B**) Heat map showing the Pearson correlation between the 22 samples. Warm colours (yellow) show highest correlation. The distinct early and late HOC phases are evident (see Figure 4). (**C**) Counts tables were generated for all *S. cerevisiae* genes and for each gene, a log_2_ fold-change vs median transformation was applied. Gap-Statistic analysis was applied to the transformed RNASeq data to determine the correct number of clusters into which to divide genes using the k-means++ algorithm. Three clusters were chosen (based on the NETSeq analysis), although no clustering is indicated by the Gap-statistic. (**D**) The transformed data was divided into three clusters using the k-means++ algorithm and plotted over time alongside all genes. (**E**) The RNASeq counts tables were normalized to gene-length, then log_2_(x+1) transformed and plotted over time and finally divided into octiles based on median expression level. Mean expression over time was plotted for each octile. Profiles were colour-coded based on cluster membership (red – HOC; blue – LOC, tan – NC). (**F**) Gap-Statistic analysis was applied to the transformed RNASeq data to determine the correct number of clusters in which to subdivide genes using the k-means++ algorithm after dividing the data into the three NETSeq clusters (NC-T, LOC-T and HOC-T). The Gap statistic indicates that the HOC-T cluster should be divided into three sub clusters using the RNASeq data whereas LOC-T and NC-T should not be subclustered. (**G**) The 3 HOC-T subclusters of the LFCM transformed RNASeq data were plotted together with the equivalent NETSeq subclusters over time. Analysis of genes in each subcluster show enrichment for those involved in protein synthesis: ribosomal protein genes and two groups of genes whose products are involved in ribosome biogenesis (RiBis) and show the differences in the RNASeq and NETSeq profiles for these gene groups.

**Figure EV3.**
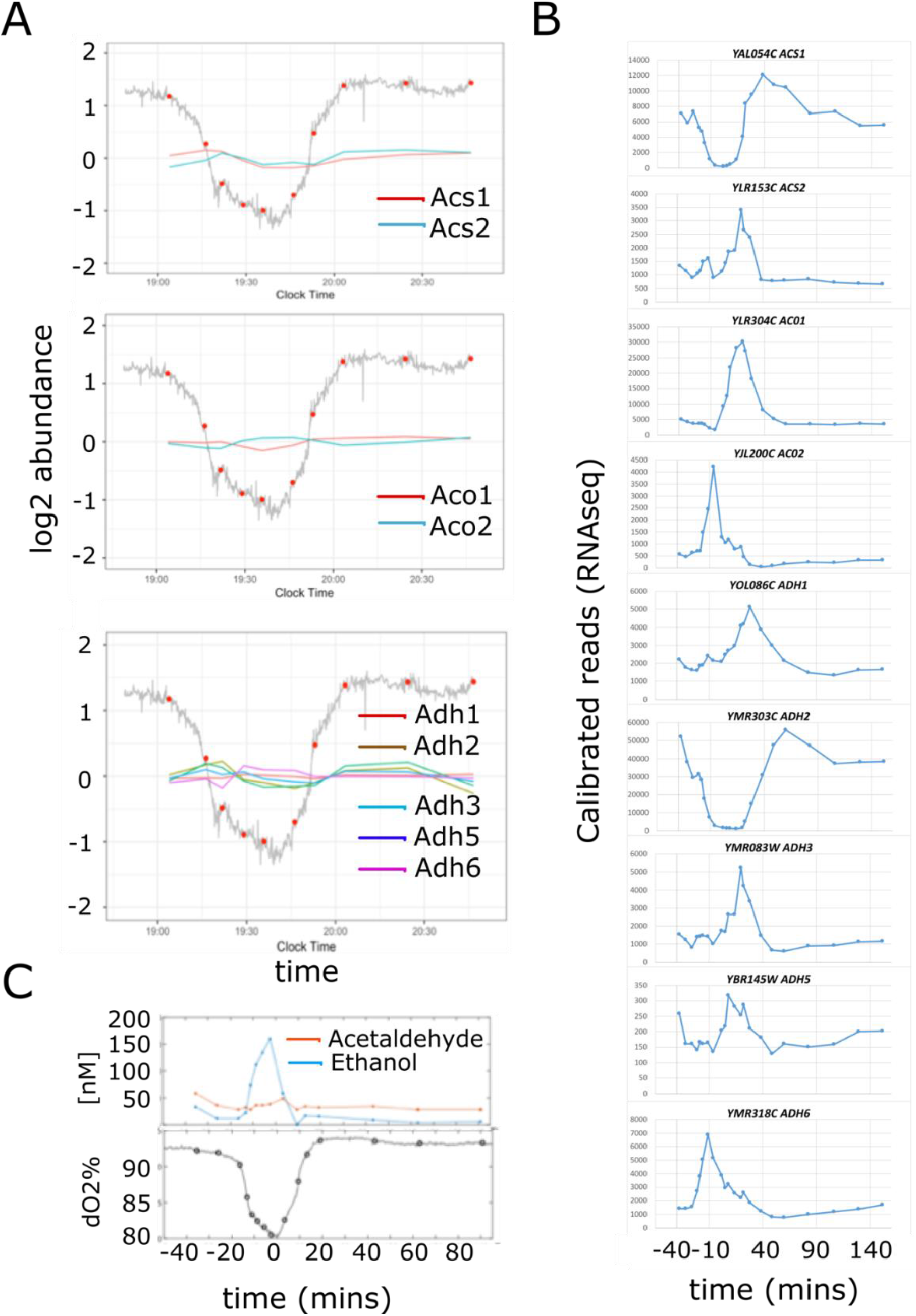
Profiles of selected metabolic enzymes and their transcripts during the YMC. (**A**) log2 abundance for selected metabolic enzymes plotted onto the oxygen consumption profiles with sampling times indicated. (**B**) Calibrated read counts for selected stable transcript levels (RNASeq) over the YMC. Sampling times are shown in Figure EV2A. (**C**) Levels of ethanol and acetaldehyde in the fermenter [nM] over time at the sampling times indicated on the dissolved oxygen plot. Ethanol is rapidly metabolized during the late HOC phase but levels of all 5 alcohol dehydrogenase *(Adh1-6)* enzymes remain largely stable throughout the cycle.

**Figure EV3 Appendix 1:**
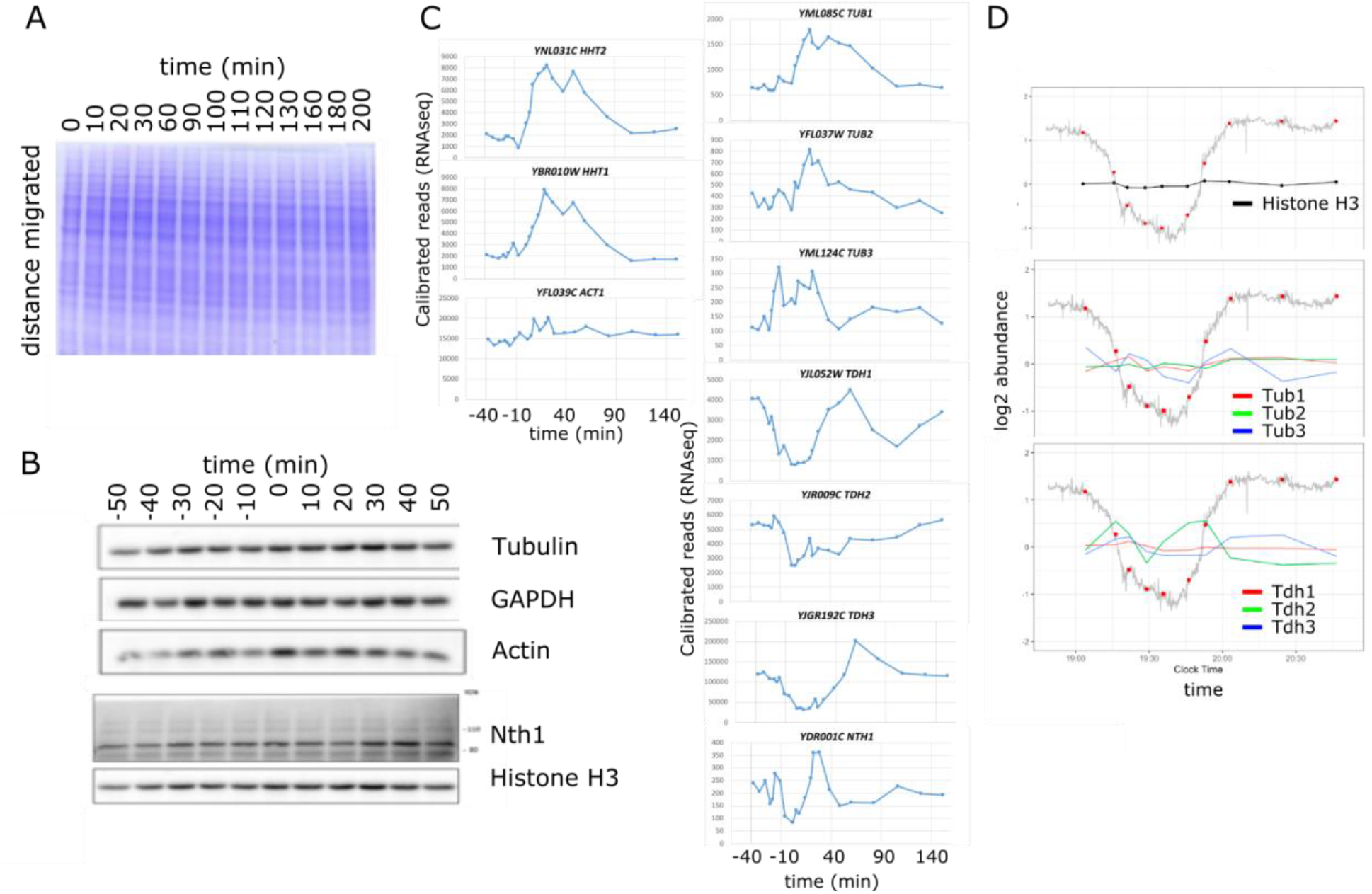
Behaviour of selected proteins and transcripts during the YMC. (**A**) Levels of total proteins remain constant throughout the YMC. Coomassie stain of total proteins extracts after separation by SDS PAGE. Time points are relative to the point of maximum oxygen-consumption. (**B, C**) Western blots (**B**) showing levels of selected proteins (tubulin, glyceraldehyde 3-phosphate dehydrogenase, actin, 13myc-tagged neutral trehalase (Nth1) and histone H3) during the YMC and the profile for the steady state levels of mRNA transcripts (**C**) encoding these proteins. (**D**) Levels of the Tubulin (Tub1-3), Glyceraldehyde 3-phosphate dehydrogenase proteins (Tdh1-3) and histone H3 proteins determined by TMT-labelled proteomics during the YMC. Although the majority of transcripts cycle (with the exception of the *ACT1* transcript), levels of the encoded proteins remain constant.

**Figure EV4.**
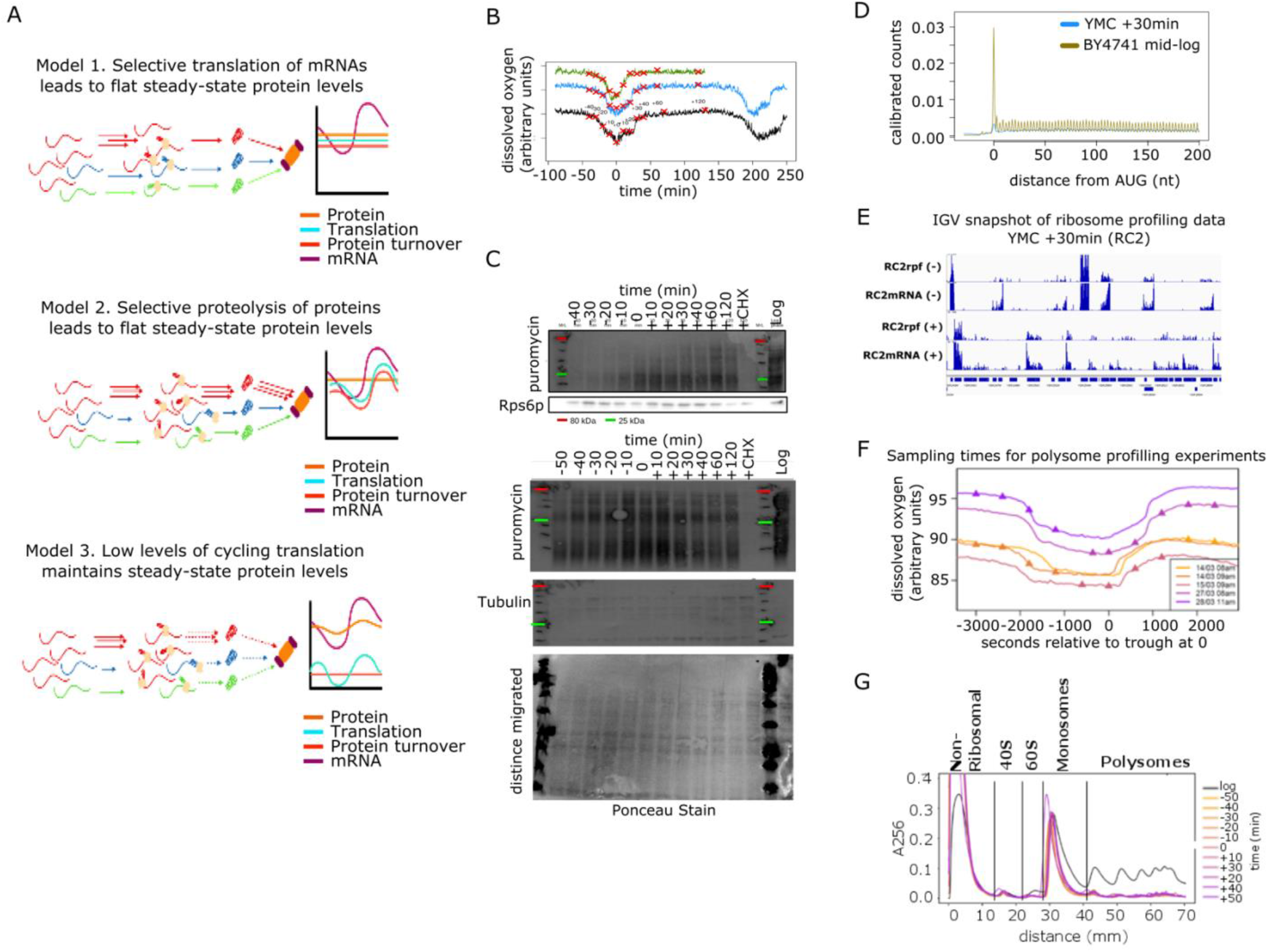
Translation cycles during the YMC. (**A**) Three models for how the proteome may be buffered against transcriptional changes. (Topmost) Translation rates are constant. Despite the transcriptome cycling, protein synthesis rates for individual genes are constitutive, meaning that protein levels too are constant. (Central) Perfect coordination of synthesis and degradation. Cycling transcripts lead to cycling protein synthesis rates. However, protein degradation rates also cycle which cancels out any changes in levels. (Bottom) Slow protein turnover minimizes the impact of cycling transcripts. Cycling of transcripts leads to cycling of protein synthesis rates. However, only a minimal amount of protein is synthesized and degraded each cycle, meaning that the amplitude of cycling protein levels is negligible. **(B)** Sampling times plotted on dissolved oxygen profiles for three experiments to assess levels of protein synthesis during the YMC using the puromycylation assay. (**C**) Representative western blots of cell extracts probed with anti-puromycin antibodies in total protein extracts after a 5-minute pulse label with puromycin at each time point shown. The top blot is shown relative to levels of Rps6 protein, the bottom blot relative to tubulin levels and a Ponceau stain showing total proteins levels on the blot. At +120 mins relative to the highest point of oxygen consumption cells were treated with or without cycloheximide to inhibit protein synthesis to control for background levels and levels in cells grown in exponential growth (log) are shown for comparison. (**D**) Metagene showing ribosomal occupancy over the first 200nt of transcripts in the YMC sampled at +30 mins relative to the point of highest oxygen consumption and in mid-log cells. (**E**) IGV (Thorvaldsdottir et al., 2013) snap-shot of ribosome profiling data at t=+30 on the Watson (+) and Crick (−) strands over a region of the yeast genome. Ribosome protected fragments (rpf) extracted from the ribosome profiling data set (RC2rpf) compared to the total levels of mRNA at the same region (RC2mRNA). This image reveals that the ribosome protected fragments extend over the whole ORF on each mRNA, although the density of ribosome protected fragments may vary on individual mRNAs. For details analysis of this data see (Feltham, 2018). (**F**) Sampling times for polysome profiling experiments. Samples were taken from 5 separate cycles at the time points shown relative to t=0 at the lowest point on the dissolved oxygen trace. Together these represent n=2 for 10 timepoints through two complete cycles of the YMC. (**G**) Polysome profiles at the times shown in the YMC relative to the point of highest oxygen consumption and from a mid-log culture in exponential growth (black line) focusing in on the difference in the non-ribosomally associated RNA for the midlog culture and the relative occupancy by monosomes for the YMC samples. Sampling times throughout the YMC are the same as in Figure 3C.

**Figure EV5.**
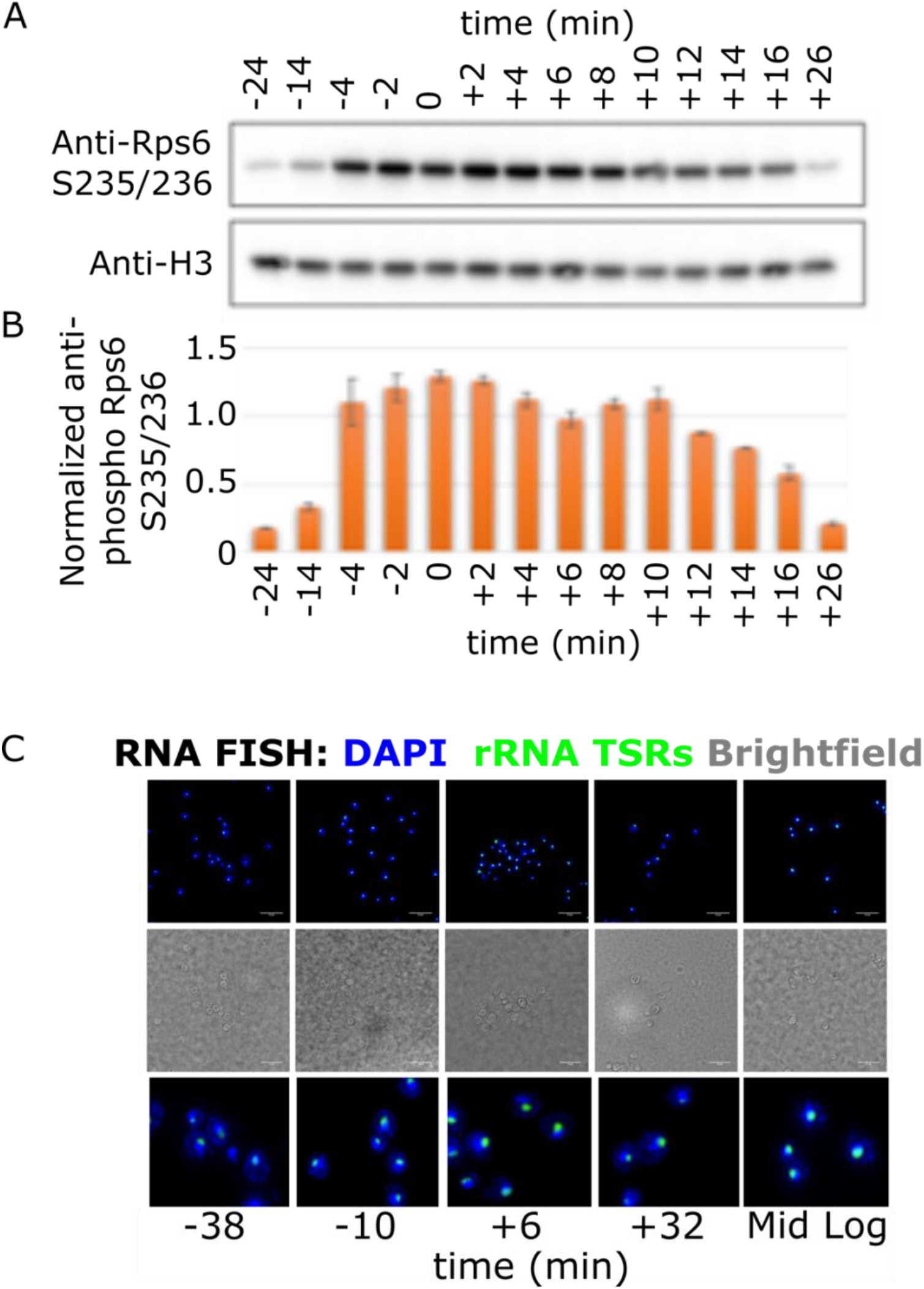
Ribosome assembly cycles during the YMC. **(A)** Western blots showing the levels of doubly-phosphorylated Rps6 at S235/S236 and histone H3, as a loading control, at 14 time-points through the YMC. (**B**) Normalisation of Rps6 S235/S236 phosphorylation to histone H3 levels (see Figure EV5 appendix 1 for 2 repeats). (**C**) Representative rRNA FISH microscopy images of samples taken during the LOC (−38, +32) vs HOC (−10,+6) of the YMC and in the mid-logarithmic phase (Mid Log). Images of the same field, using fluorescence (top and bottom cropped and enlarged) and brightfield (middle). Fluorescent labeling; DNA is stained with DAPI (blue) and transcribed spacers regions of the 35S ribosomal RNA are probed with DNA probes labelled with Quasar 570 in green.

**Figure EV5 Appendix 1.**
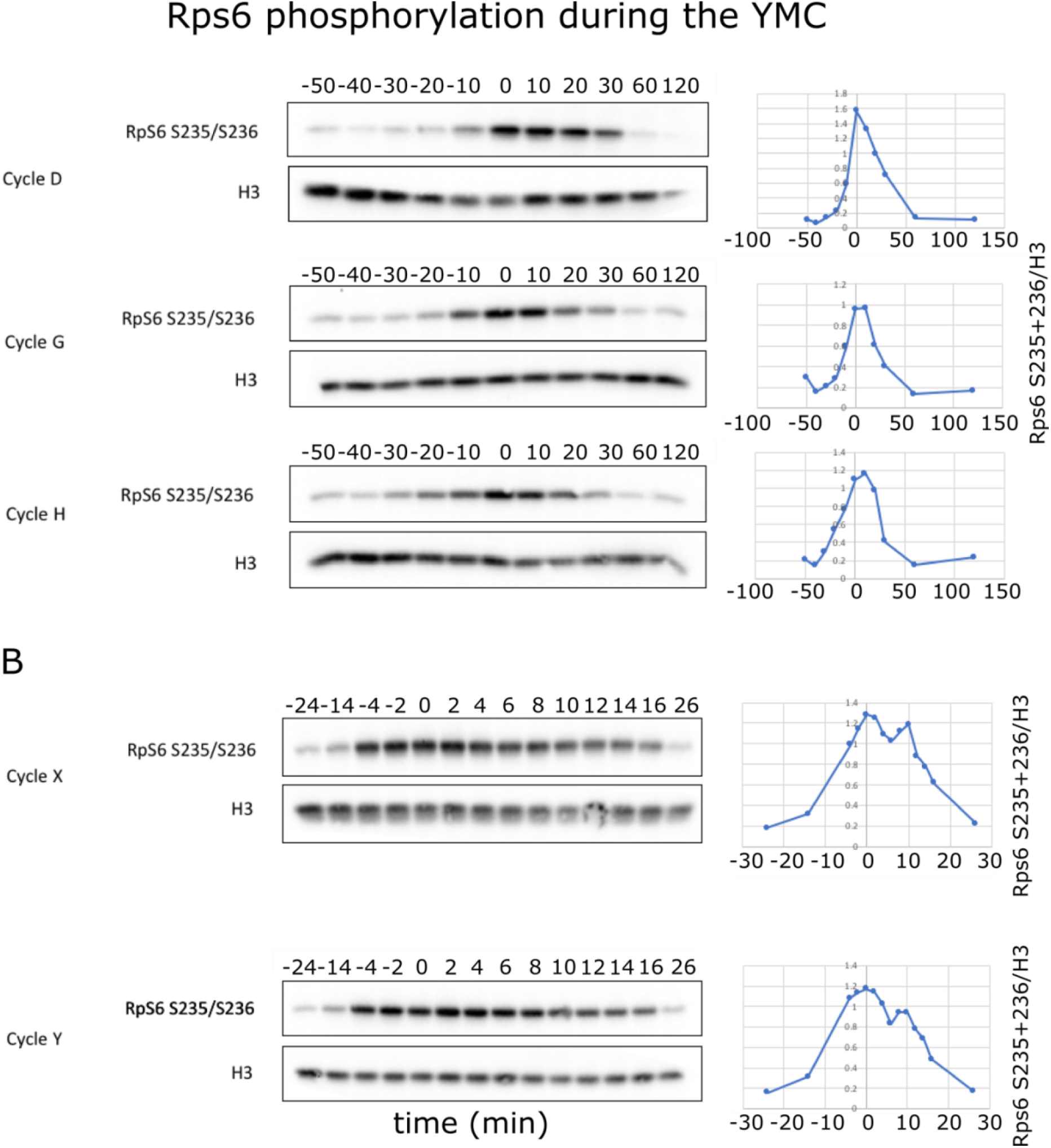
Rps6 phosphorylation cycles during the YMC. **(A)** Western blots showing the levels of doubly-phosphorylated Rps6 at S235/S236 and histone H3, as a loading control, at 11 time-points of three different cycles through the YMC. For each blot, levels of Rps6p protein relative to histone H3 are shown. This data is used in Figure 4B. (**B**) Western blots showing the levels of doubly-phosphorylated Rps6 at S235/S236 and histone H3, as a loading control, at 14 time-points of two different cycles through the YMC. For each blot levels of Rps6p protein to histone H3 are shown.

**Figure EV5 Appendix 2:**
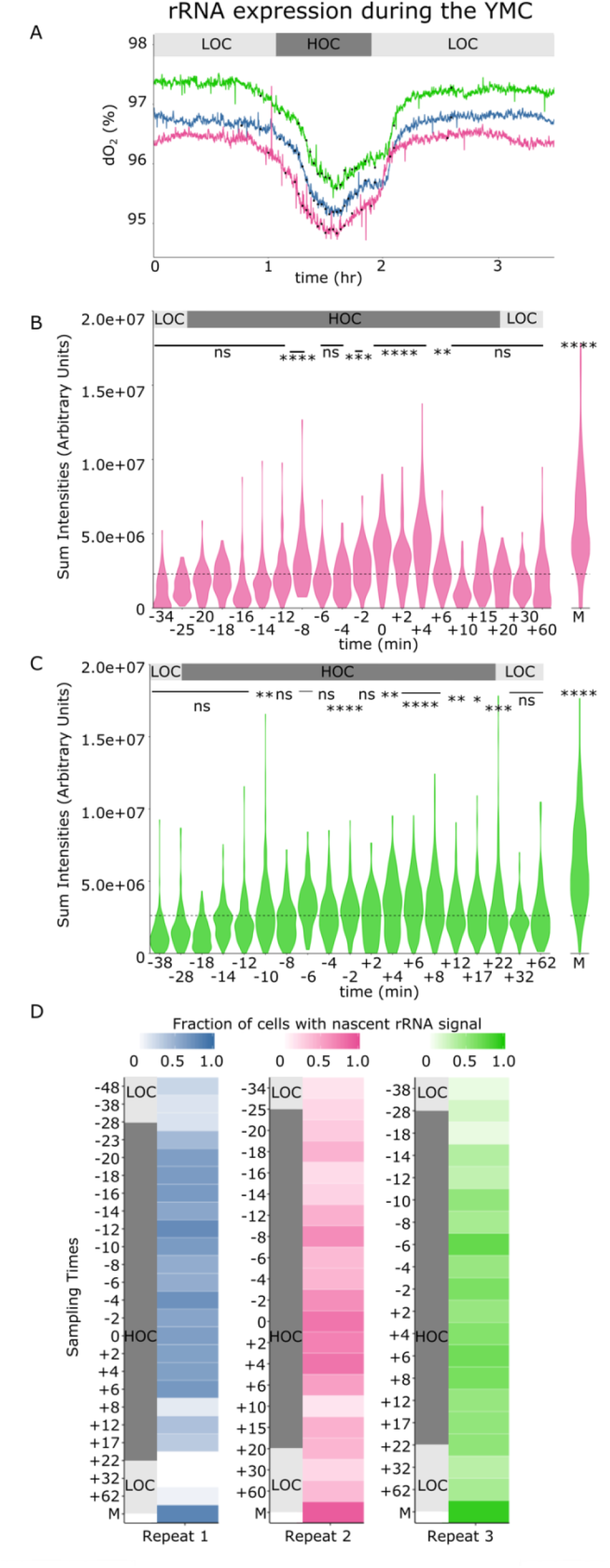
rRNA expression during the YMC. **(A)** 20 sampling points for each biological replicate used for the rRNA FISH experiment shown relative to the dO_2_ % levels. The upper background colour indicates the phase of the YMC; light grey = Low Oxygen Consumption, grey = High Oxygen Consumption. **(B,C)** Violin plots showing the sum of the rRNA intensities determined by RNA FISH to spacer regions, allowing assessment of rates of transcription before processing. Levels of rRNA production peak in the HOC phase of the cycle. Levels are substantially lower than in exponentially growing cultures. The sampling time points are shown in minutes relative to the point of lowest dO_2_ % in the YMC, displayed as “0”. “M” indicates the sample taken in the middle of the logarithmic phase. The upper background colour indicates the phase of the YMC; light grey = Low Oxygen Consumption, grey = High Oxygen Consumption. The dashed line represents the base mean of all the time points sum intensities excluding the sample taken in the middle of the logarithmic phase. Asterisks indicate time points for which rRNA sum intensities are significantly greater than the base mean (p-value<0.05). P-values were calculated by Wilcoxon test. Second and third repeat of three biological replicates. **(D)** Heat maps presenting the fraction of cells showing greater rRNA sum intensities than the base mean of all the time points sum intensities calculated without taking into account the sample taken in the middle of the logarithmic phase over the total number of cells detected. The proportion of cells expressing high rRNA levels increases during the HOC phase of the YMC. Each row of the heat maps represents a sampling time point in minutes relative to the point of lowest dO_2_ % in the YMC, displayed as “0”. “M” indicates the sample taken in the middle of the logarithmic phase. The background colour on the left indicates the phase of the YMC; light grey = Low Oxygen Consumption, grey = High Oxygen Consumption. Each heat map corresponds to a biological replicate (R1, R2, R3). During the late LOC phase only 15% of the cells are expressing high rRNA levels. This fraction increases when entering the HOC phase with a peak of the rRNA levels around the middle (from −20 to −6 depending on the biological replicate) where more than 65% cells are expressing high rRNA levels, showing an overall 4-fold change in the number of cells expressing high rRNA levels. Finally this percentage of cells decreases during the later HOC phase and returns to an average of 30% of cells expressing high rRNA levels when entering the early LOC phase. High is higher than the base mean. Thus not only the overall rRNA levels increases as oxygen consumption increased, but there is a shift in the cell population towards expressing more rRNA. However, a few cells per set do not express rRNA. The images suggest that the majority of the non-expressing cells are newly formed cells, budded cells. Taken together with the data in Figure 4A,B, Figure EV5, and Figure EV5 appendix 1, these results suggest that ribosome assembly is coordinated during the YMC.

**Figure EV5 Appendix 3:**
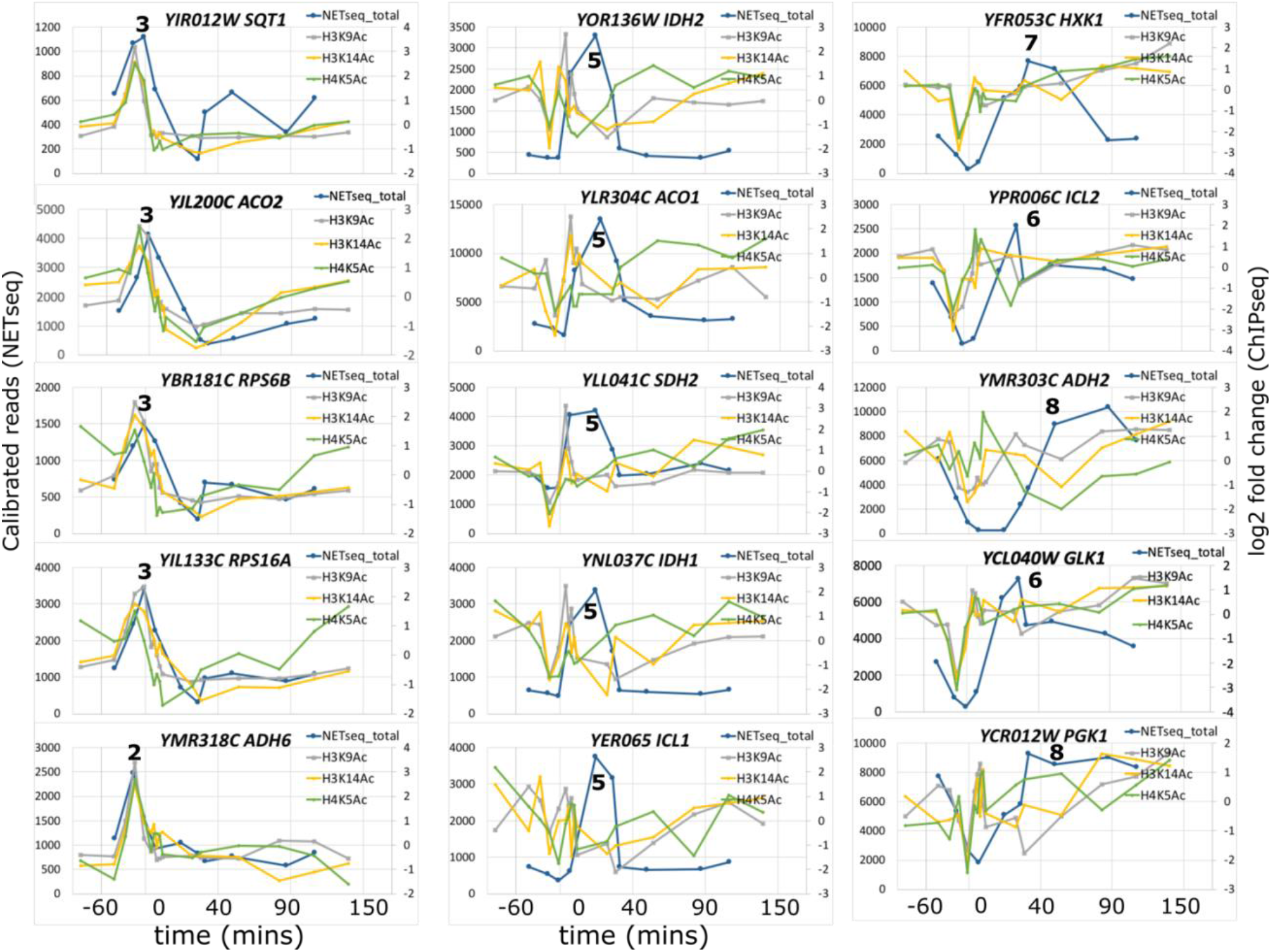
Examples showing the relationship between acetylation and nascent transcript production in the YMC. Plots showing log_2_-fold changes in levels of histone acetylation (H3K9ac, H3K14ac and H4K5ac) at gene promoters compared to the levels of nascent transcripts reads over time where 0 is the point of lowest dissolved oxygen. The three columns show selected genes whose transcription peaks in early HOC (left) later HOC (middle) or during LOC (right) and the associated changes in histone acetylation. The number indicated the sample point for reference, with details sampling times shown in EV Table 1, with 10 time points for NET-seq and 16 time points for ChIP-seq.

**EV Table 1.** Excel file with source data for transcriptomics (NETSeq, RNA-seq), ChIP-seq (histone modifications), metabolites and a macro to allow parameters for each gene to be displayed over time.

